# The PST repeat region of MDC1 is a tunable multivalent chromatin tethering domain

**DOI:** 10.1101/2025.01.10.632395

**Authors:** Joshua R. Heyza, Maria Mikhova, Gloria I. Perez, David G. Broadbent, Jens C. Schmidt

## Abstract

DNA double strand breaks (DSBs) are widely considered the most cytotoxic DNA lesions occurring in cells because they physically disrupt the connectivity of the DNA double helix. Homologous recombination (HR) is a high-fidelity DSB repair pathway that copies the sequence spanning the DNA break from a homologous template, most commonly the sister chromatid. How both DNA ends, and the sister chromatid are held in close proximity during HR is unknown. Here we demonstrate that the PST repeat region of MDC1 is a mutlivalent nucleosome binding domain, sufficient to tether chromatin in multiple contexts. In mitotic cells the affinity of the PST repeats for chromatin is downregulated by phosphorylation to prevent chromosome missegregation, while still contributing to DNA break tethering by the MDC1-TOPBP1-CIP2A complex. In interphase, the PST repeat region is critical for RAD51 focus formation but not the recruitment of 53BP1 to DNA breaks, consistent with a chromatin tethering function. In total, this work demonstrates that the PST repeat region of MDC1 is a multivalent chromatin binding domain with tunable affinity that contributes to DNA break tethering during HR and in mitosis.

## INTRODUCTION

DNA double-strand breaks (DSB) are widely considered to be the most toxic genomic lesion. DSBs can arise in all cell-cycle phases and are induced both as part of intrinsic cellular processes and as a consequence of exposure to endogenous and exogenous chemicals ^1^. These lesions pose significant threats to genome stability because the failure to repair DSBs can lead to loss of genetic information, insertions and deletions, or translocations which can result in the development of a wide array of diseases including immunodeficiencies and cancers ^2–4^. Induction of DSBs and targeting of pathways that repair them are widely used approaches for cancer treatment ^5,6^. In addition, specific DSB formation by Cas enzymes and their controlled repair are critical steps in genome editing, which has tremendous therapeutic potential ^7^. Comprehensively defining the molecular mechanisms underlying DSB repair therefore has significant implications for human health.

Three major kinases mediate the cellular response to DSBs, including DNA-dependent Protein Kinase catalytic subunit (DNA-PKcs), Ataxia Telangiectasia Mutated (ATM), and ATM and Rad3-related (ATR) ^8^. These kinases initiate a chromatin signaling cascade at DSBs by phosphorylating histone H2AX at S139 (γH2AX) ^8^. γH2AX is bound by the adaptor protein, Mediator of DNA damage checkpoint 1 (MDC1) through its C-terminal BRCT domain ^9,10^. MDC1 is a large protein consisting of 2089 amino acids with several well characterized functional domains ^11^. MDC1 carries out several important functions in genome maintenance throughout the cell cycle. At chromosomal DSBs in interphase, MDC1 promotes high-fidelity homologous recombination (HR) and error-prone non-homologous end joining (NHEJ) repair by participating in signaling cascades that recruit BRCA1 and 53BP1 to DNA breaks ^9,10,12–17^. This is accomplished by a direct interaction between MDC1 and several repair factors including the E3 ubiquitin ligase RNF8, which binds to a cluster of TQXF motifs that are phosphorylated by ATM after DSB induction ^15^. This interaction initiates a series of chromatin ubiquitination events where RNF8 poly-ubiquitinates histone H1, creating a docking site for a second E3 ubiquitin ligase RNF168 which facilitates BRCA1 and 53BP1 localization to sites of DNA damage by depositing ubiquitin moieties on H2A at K13 and K15 ^17–20^. BRCA1 and 53BP1 in turn carry out critical effector functions in the HR and NHEJ repair pathways, respectively ^21–28^.

In mitosis cells are particularly vulnerable to DSBs, because the presence of unrepaired DNA breaks can lead to a loss or missegregation of the chromosome arm not connected to the centromere ^29^. As a repair pathway of last resort, recent work has shown that alternative end joining mediated by polymerase theta is activated in mitosis via the adapter protein RHINO ^30^. In contast, during cell division HR and 53BP1-dependent end joining are suppressed by inhibiting the RNF8-MDC1 interaction via CDK1-mediated phosphorylation of RNF8 at T198 and also by phosphorylation of 53BP1 ^31^. Nevertheless, MDC1 still binds to mitotic DSBs marked by γH2AX where phosphorylation of MDC1 at S168 and S196 by CK2 enables an interaction with TOPBP1 and the formation of the MDC1-TOPBP1-CIP2A complex that tethers broken DNA ends together throughout mitosis and into G1-phase where they are repaired by 53BP1-dependent end joining ^32,33^. Additionally, the MDC1-TOPBP1-CIP2A axis has been implicated in mitotic tethering of shattered micronuclear chromosome fragments ^34,35^.

Because DSBs lead to a loss of connectivity of the broken DNA ends their effective and rapid tethering is a critical step in DNA break repair. In classical NHEJ, DNA ends are rapidly tethered by the Ku70/80 hetero-dimer and DNA-PKcs, and 53BP1 plays an important role in stabilizing DSB ends in NHEJ downstream of MDC1 ^36,37^. In the context of HR, it is unclear how DNA ends are held together. HR is a prolonged and complex process that takes many steps for accurate repair of a DSB ^38^. In addition to requiring the tethering of the two ends of the DNA break, HR also relies on access to the sister chromatid to copy the sequence spanning the DSB. How the DNA ends and the sister chromatin are held in close proximity for an extended period of time during HR is unknown.

In addition to several well-characterized domains and protein-protein interaction sites, MDC1 also contains a large, intrinsically disordered region referred to as the PST (Proline-Serine-Threonine) repeat domain whose function remains poorly understood. The PST domain first arose in mammalian MDC1 and is defined by a largely conserved 41 amino acid sequence repeated in tandem with human MDC1 containing 13 repeats ^39,40^. The PST repeat region has been implicated in DSB repair and its deletion impairs BRCA1 focus formation and HR without impacting 53BP1 focus formation ^13^. However, the precise function of the PST domain in HR remains unclear. Interactions between MDC1’s PST domain and DNA-PKcs, Chk1, FBXW7, Rag1, and pre-rRNA have been reported and the PST domain has been linked to Chk1 activation and ubiquitin-dependent regulation of MDC1 protein stability ^41–44^. Using live-cell single-molecule imaging and biochemical approaches, work from our group and Salguero *et al.* recently revealed that MDC1 is constitutively associated with chromatin in a DNA damage-independent manner in interphase via an interaction of the PST repeat domain with the nucleosome acidic patch ^45,46^.

In this study, we demonstrate that the PST repeat domain of MDC1 is a multivalent chromatin binding domain sufficient for DNA tethering. We observe that the constitutive chromatin association of MDC1 mediated by the PST repeat region is suppressed in mitosis by CDK1-mediated phosphorylation of 21 Threonine-Proline (TP) motifs in the PST repeats. Using custom phospho-specific antibodies we demonstrate that PST repeat phosphorylation occurs upon nuclear envelope breakdown and phosphorylated MDC1 foci at mitotic DSBs persists into early G1 phase. While mitotic phosphorylation of the PST repeats does not impact MDC1 focus formation or recruitment of TOPBP1/CIP2A, we find that loss of the PST domain impairs the recruitment of the MDC1-TOPBP1-CIP2A complex to DNA breaks in mitosis. Importantly, preventing mitotic phosphorylation of the PST repeats leads to accumulation of MDC1 on mitotic chromatin, “gluing” condensed mitotic chromosomes together and preventing their separation during anaphase. In addition, we show that local accumulation of the PST repeat region alone on a single mitotic chromsomes is sufficient to trigger chromosome missegregation. In interphase cells the PST repeat region is dispensable for 53BP1 recruitment but required for the loading of RAD51 to mediate HR. Our results demonstrate that this phenotype is unlikely caused by a reduction in ubiquitin signaling and that altering the affinity of the PST repeat region for chromatin impairs HR. We therefore propose a model in which accumulation of MDC1 at DSBs is critical to crosslink the chromatin surrounding the DNA breaks to ensure DNA ends and the sister chromatid are maintained in close proximity to facilitate efficient DNA break repair by homologous recombination.

## RESULTS

### MDC1 is displaced from chromatin during mitosis

In previous work using live-cell single-molecule imaging we demonstrated that HaloTagged MDC1 expressed from its endogenous locus in U2OS cells is constitutively associated with chromatin in interphase cells (Fig. 1A,B) ^45^. To characterize the properties of MDC1 in a second cell line, we introduced the HaloTag into the endogenous *MDC1* locus in RPE1 cells (Fig. 1C). During our single-molecule imaging experiments in U2OS cells we observed rare cells that displayed high Halo-MDC1 mobility, which appeared to be undergoing cell division. To determine the dynamics of Halo-MDC1 diffusion during mitosis, we synchronized cells in M-phase using nocodazole and carried out live cell single-molecule imaging experiments using highly inclined laminated optical sheet microscopy (Fig. 1A, Movie S1-6) ^47,48^. Single-particle tracking and data analysis using the SPOT ON tool demonstrated that in mitotic U2OS and RPE1 cells the fraction of static Halo-MDC1 molecules, which represents MDC1 bound to chromatin, was ∼20%, while ∼60% and ∼75% of Halo-MDC1 molecules were immobile in interphase U2OS and RPE1 cells, respectively (Fig. 1D, Fig. S1A-C) ^49^. In addition, the diffusion coefficients of both mobile and static Halo-MDC1 molecules were increased in mitotic compared to interphase cells in both U2OS and RPE1 cells (Fig. 1F-I). These observations demonstrate that MDC1 is displaced from chromatin when cells enter mitosis.

**Figure 1.**
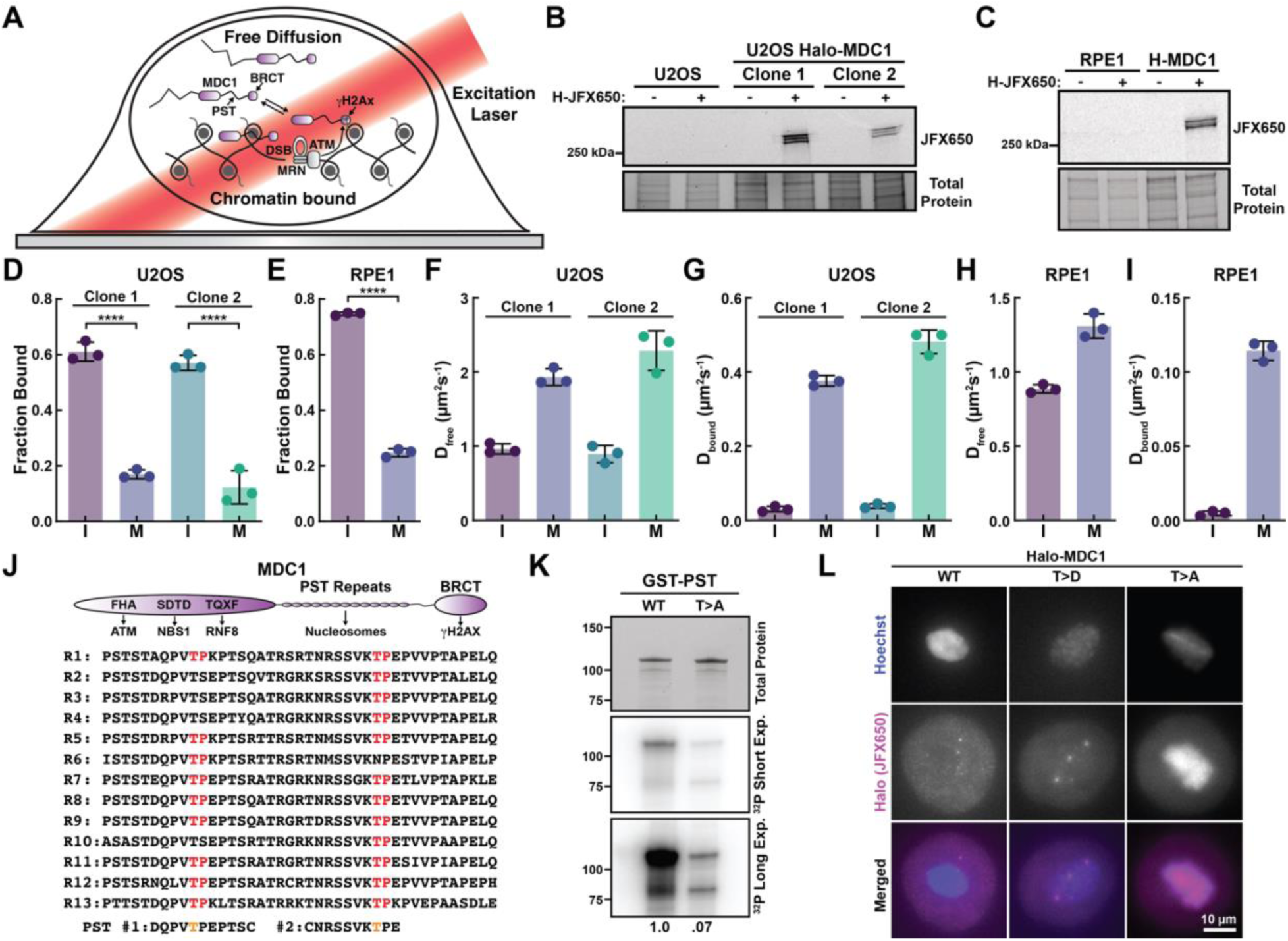
Phosphorylation of the PST repeat region removes MDC1 from chromatin in mitosis. **(A)** Model of MDC1 recruitment to chromatin and HILO microscopy for single-molecule imaging. **(B-C)** Fluorescence gels of cell lysates from **(B)** U2OS and **(C)** RPE1 expressing HaloTagged MDC1 from its endogenous locus labeled with JFX650 HaloTag-ligand. **(D-E)** Static fraction of Halo-MDC1 molecules in interphase (I) and mitosis (M) derived from single-particle tracking of MDC1 in **(D)** U2OS and **(E)** RPE1 cells (N = 3 biological replicates, ≥ 20 cells per replicate, Mean ± SD, two-tailed T-test). **(F-G)** Diffusion coefficient of **(F)** mobile and **(G)** static Halo-MDC1 molecules in interphase (I) and mitosis (M) derived from single-particle tracking of MDC1 in U2OS cells (N = 3 biological replicates, ≥ 20 cells per replicate, Mean ± S.D., two-tailed T-test). **(H-I)** Diffusion coefficient of **(H)** mobile and **(I)** static Halo-MDC1 molecules in interphase (I) and mitosis (M) derived from single-particle tracking of MDC1 in RPE1 cells (N = 3 biological replicates, ≥ 20 cells per replicate, Mean ± S.D., two-tailed T-test). **(J)** Domain organization of MDC1, sequence alignment of the thirteen 41 amino acid PST repeats, and consensus sequence surrounding the two conserved TP sites within the PST repeats. **(K)** Phosphorescence imaging and protein staining of and SDS PAGE gel loaded with samples of the wild type (WT) and non-phosphorylatable (T>A) GST-tagged PST repeat domain treated with CDK1-Cyclin B and ATP γ-^32^P. **(L)** Images of mitotic U2OS cells with MDC1 knock-out (arrested with nocodazole) transiently expressing HaloTagged wildtype MDC1 (WT) and MDC1 variants with phospo-mimicking (T>D) and non-phosphorylatable (T>A) mutations in the TP sites within the PST repeat region.

### Phosphorylation of its PST repeat region by CDK1-Cyclin B displaces MDC1 from chromatin during mitosis

To define the molecular mechanism by which MDC1 is displaced from chromatin during mitosis, we focused on the PST repeat region of MDC1, which mediates its constitutive chromatin association in interphase cells ^45,46^. The human PST-region is composed of 13 imperfect 41 amino acid repeats (Fig. 1J). Analysis of MDC1 using the phosphosite.org data base revealed that the PST repeats contain two threonine-proline (TP) sites, which have been shown to be phosphorylated in multiple high-throughput phospho-proteomic studies ^50–53^. TP di-peptides are consensus sites for the mitotic CDK1-Cyclin B ^54^. To test whether CDK1-Cyclin B can phosphorylate the PST repeat region of MDC1, we purified the GST-tagged wildtype PST repeat region of MDC1 and a variant of the PST repeat region in which all threonine residues within the TP sites were mutated to alanine (T>A, Fig. S1D). After incubation with CDK1-Cyclin B and ATP γ-^32^P, the wildtype PST repeat region was robustly phosphorylated by CDK1-Cyclin B, while phosphorylation of the T>A variant was reduced greater than 10-fold (Fig. 1K), demonstrating that the TP sites within the PST repeats can be targeted by CDK1-Cyclin B. To determine whether phosphorylation of the TP sites within the PST repeat region is sufficient to displace MDC1 from chromatin during mitosis, we transiently expressed non-phosphorylatable (T>A) and phospho-mimicking (T>D) HaloTagged MDC1 variants in U2OS cells in which MDC1 had been knocked out (ΔMDC1) (Fig. S1D). Wildtype MDC1 and phospho-mimicking T>D MDC1 were not enriched on chromatin in nocodazole arrested mitotic cells (Fig. 1L). In contrast, non-phosphorylatable MDC1 was enriched on mitotic chromosomes (Fig. 1L). These results demonstrate that phosphorylation of the TP sites within the PST repeat region of MDC1 is necessary and sufficient to displace MDC1 from mitotic chromosomes.

### The number of PST repeats and their phosphorylation control the constitutive chromatin association of MDC1

To determine how the copy number of PST repeats and the phosphorylation of the TP sites in the repeats regulate the constitutive chromatin association of MDC1, we transiently expressed HaloTagged MDC1 variants in MDC1 knock-out cells and carried out live cell single-molecule imaging experiments (Fig. 2A, Fig. S1D). In interphase cells, the fraction of chromatin associated MDC1 gradually decreased with removal of PST repeats (Fig. 2B, Movie S7-14). Similarly, the diffusion coefficients of mobile and static MDC1 molecules gradually increased when more PST repeats were removed from MDC1 (Fig. 2C,D). This suggests that the contribution of each PST repeat to chromatin binding of MDC1 is comparable, rather than specific repeats mediating the association of MDC1 with nucleosomes.

**Figure 2.**
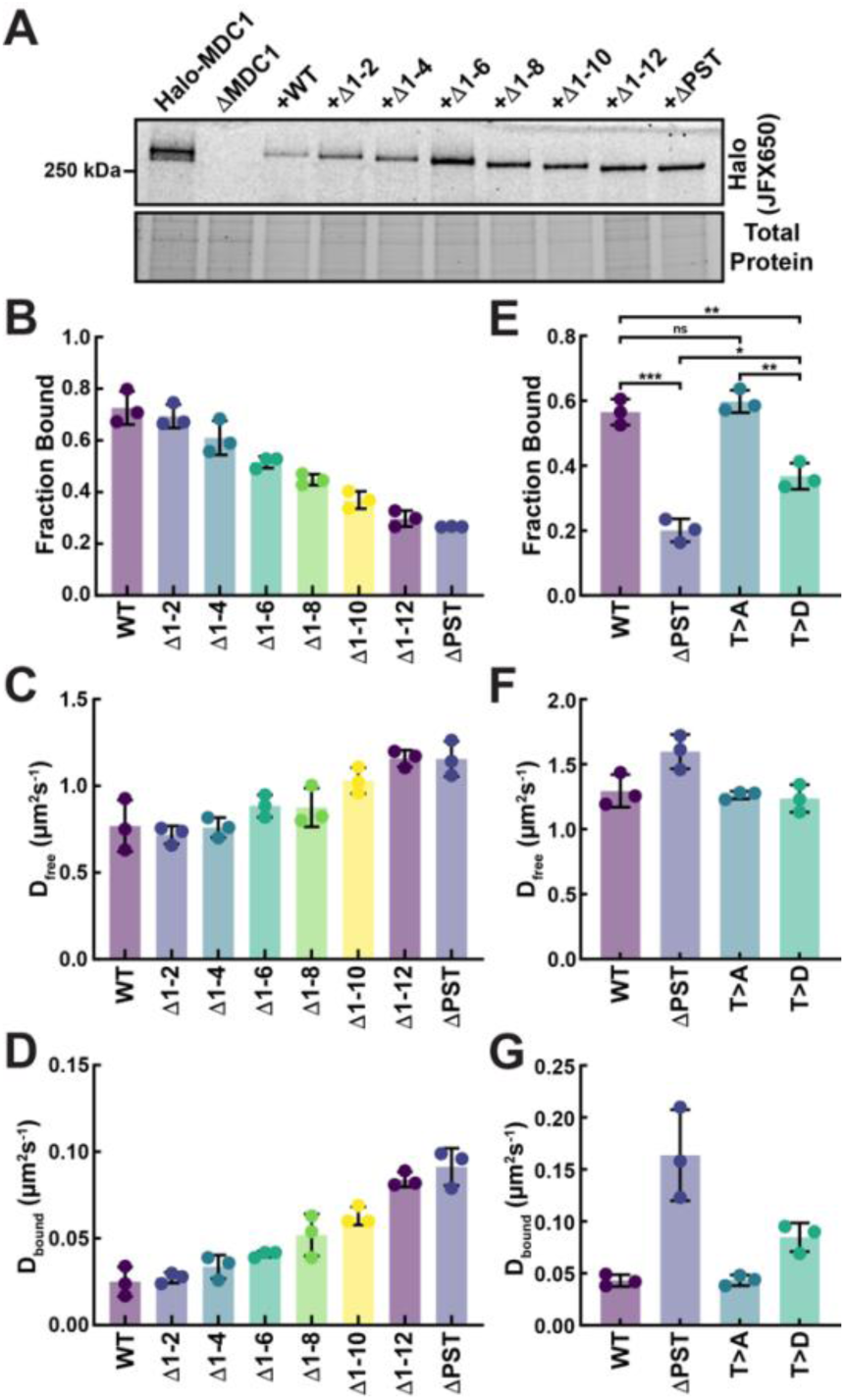
The number of the PST repeats and TP phosphorylation regulates the chromatin association of MDC1. **(A)** Fluorescence gel of cell lysates of U2OS ΔMDC1 cells transiently expressing HaloTagged (JFX650) wildtype MDC1 and truncations of the PST repeats. **(B-D)** Single particle tracking analysis of HaloTagged wildtype MDC1 and truncations of the PST repeats in interphase U2OS ΔMDC1 cells including the **(B)** fraction of static particles and the diffusion coefficients of **(C)** mobile and **(D)** static MDC1 molecules (N = 3 biological replicates, ≥ 20 cells per replicate, Mean ± S.D.). **(E-G)** Single particle tracking analysis of HaloTagged wildtype MDC1 (WT), MDC1 with a deletion of the PST repeat region (ΔPST) and MDC1 variants with phospo-mimicking (T>D) and non-phosphorylatable (T>A) mutations in the TP sites within the PST repeat region in interphase U2OS ΔMDC1 cells including the **(E)** fraction of static particles and the diffusion coefficients of **(F)** mobile and **(G)** static MDC1 molecules (N = 3 biological replicates, ≥ 20 cells per replicate, Mean ± S.D.).

The dynamic properties of non-phosphorylatable (T>A) MDC1 were indistinguishable from wildtype MDC1 (Fig. 2E-G, Movie S15-18), suggesting that the TP sites in MDC1 are not phosphorylated in interphase cells. In contrast, MDC1 with phospho-mimicking (T>D) TP sites in the PST repeat region had a reduced fraction of chromatin bound molecules and an increased diffusion coefficient of static MDC1 molecules (Fig. 2E-G). Both the reduction in the static fraction and increase in the diffusion coefficient of chromatin associated MDC1 T>D molecules was intermediate between wildtype MDC1 and MDC1 with a complete deletion of the PST repeat region (Fig. 2E-G). This suggests that MDC1 with phospho-mimicking (T>D) TP sites in the PST repeat region has a reduced affinity for chromatin, but it retains some ability to bind to nucleosomes.

In total, these observations demonstrate that the ability of MDC1 to associate with chromatin is regulated by the copy number of and phosphorylation state of the TP sites within the PST repeats. In addition, these observations are consistent with the PST repeat region forming multivalent interactions with chromatin, simultaneously binding to several nucleosomes.

### Phosphorylated MDC1 accumulates at DNA breaks on mitotic chromosomes

To further analyze the mitotic phosphorylation of MDC1 we raised phospho-specific antibodies against two phosphorylated peptides representing the consensus sequences for the two TP sites within the PST repeats (Fig. 1J). Both phospho-specific antibodies recognized Halo-MDC1 foci in zeocin treated mitotic but not interphase cells (Fig. 3A,B), demonstrating that phosphorylated MDC1 is recruited to sites of DNA damage in mitosis. To confirm the specificity of antibodies raised against the phosphorylated PST repeat epitopes, we knocked out MDC1 in U2OS cells and re-expressed HaloTagged MDC1 variants from a doxycycline inducible expression cassette integrated into the genome using the Sleeping Beauty transposon system (Fig. S2A) ^55^. In mitotic cells treated with zeocin, the immunofluorescence signal from the antibody raised against the first TP site co-localized the wildtype Halo-MDC1 foci, but no phospho-MDC1 foci were detected in cells expressing Halo-MDC1 with a deletion of the entire PST repeat region (ΔPST), or Halo-MDC1 with non-phosphorylatable (T>A) or phospho-mimicking (T>D) TP sites within the PST repeat region, confirming the specificity of the antibody (Fig. 3C). Importantly, Halo-MDC1 lacking the PST repeat region had greatly impaired DNA damage induced foci formation in mitotic cells treated with zeocin, while all other Halo-MDC1 variants tested (WT, T>A, T>D) formed foci (Fig. 3C). These observations demonstrate that phosphorylated MDC1 accumulates at DNA breaks in mitosis which requires the PST repeat region.

**Figure 3.**
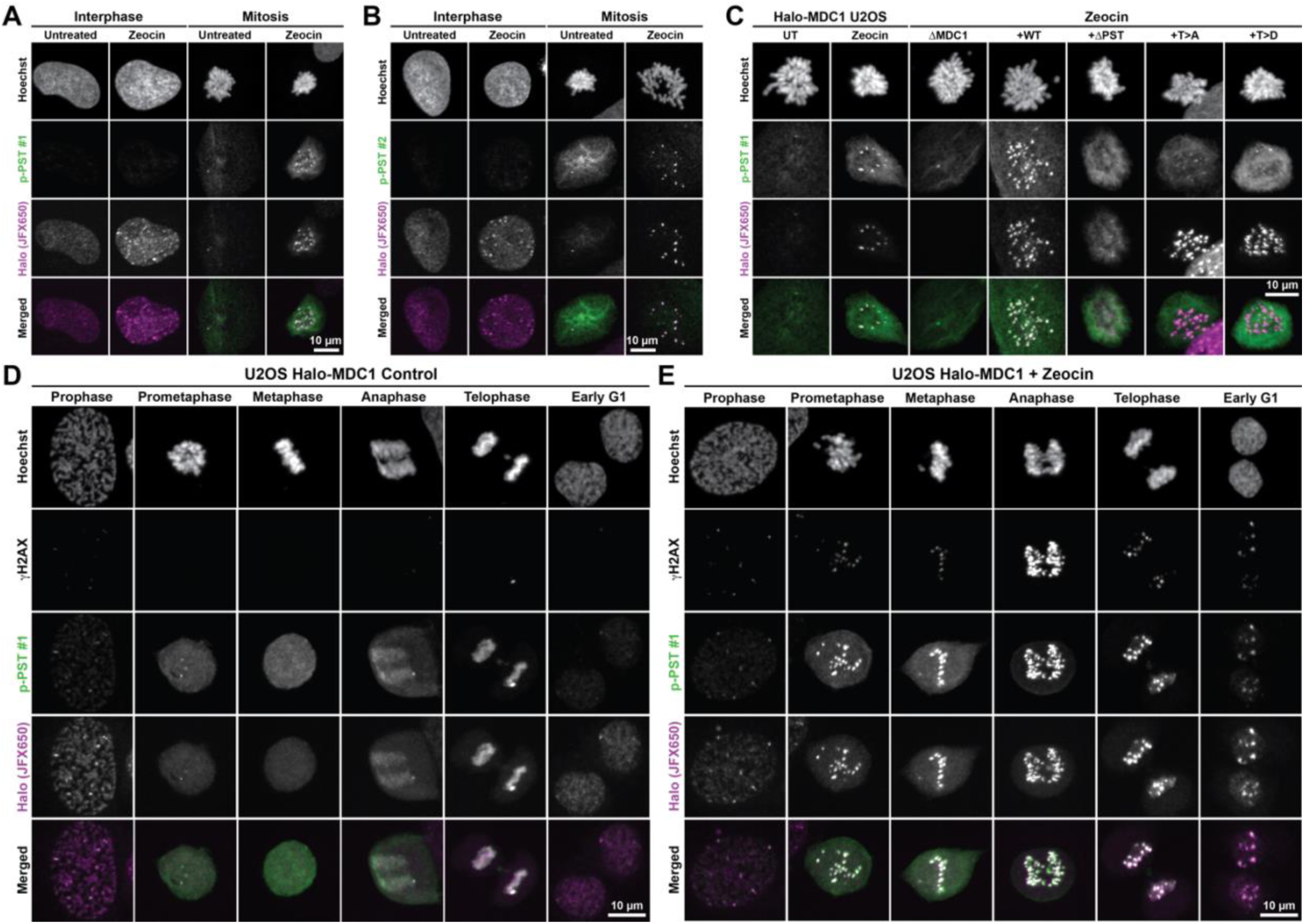
Phosphorylated MDC1 accumulates at DNA breaks in mitosis and early G1-phase. **(A-B)** Images of fixed interphase and mitotic U2OS cells expressing Halo-MDC1 (JFX650) immuno-stained with **(A)** p-PST antibody #1 or **(B)** p-PST antibody #2 and Hoechst to label DNA. **(C)** Images of fixed, mitotic U2OS cells expressing Halo-MDC1 and U2OS ΔMDC1 cells stably expressing HaloTagged MDC1 (JFX650) variants untreated or treated with zeocin. Cells were immuno-stained with p-PST antibody #1 and Hoechst to label DNA. **(D-E)** Images of fixed U2OS cells expressing Halo-MDC1 (JFX650) **(D)** untreated or **(E)** treated with zeocin to induce DNA breaks in various phases of mitosis and early G1-Phase. Cells were immuno-stained with p-PST antibody #1, a γH2AX antibody, and Hoechst to label DNA.

### Mitotic DNA breaks marked by phosphorylated MDC1 transition into 53BP1 nuclear bodies in G1

To determine the timing of MDC1 phosphorylation throughout cell division we analyzed fixed cells in different stages of mitosis. In control cells not exposed to a DNA damaging agent, phosphorylated MDC1 was diffusely localized throughout the cell during mitosis (Fig. 3D). Upon anaphase onset phosphorylated MDC1 was detected throughout cell and begun to enrich on DNA (Fig. 3D). In telophase phosphorylated MDC1 was strongly enriched on chromatin (Fig. 3D). This suggests that MDC1 dephosphorylating is initiated immediately upon anaphase onset, and MDC1 rapidly transitions to its constitutively chromatin bound state upon mitotic exit. In cells treated with zeocin to induced DSBs, phosphorylated MDC1 co-localized with DNA lesions marked by γH2AX starting upon chromosome condensation in prophase (Fig. 3E). Phosphorylated MDC1 was localized to sites of DNA damage throughout mitosis and was retained at DNA lesions into G1-phase of the next cell cycle (Fig. 3E). These results were confirmed with the phospho-specific antibody targeting the second TP site within the PST repeats (Fig. S2B,C). To determine whether DNA breaks marked by phosphorylated MDC1 transition into 53BP1 nuclear bodies after entry into G1, we carried out immunofluorescence experiments to detect 53BP1 and Halo-MDC1. As expected, 53BP1 was not detected at mitotic DSBs (Fig. S2D). Upon entry into G1-phase DNA breaks marked by γH2AX co-localized with Halo-MDC1 and 53BP1 signals (Fig. S2D). This demonstrates that DNA breaks marked by phosphorylated MDC1 in mitosis transition into 53BP1 nuclear bodies in G1-phase of the cell cycle.

### The PST repeat region is required for MDC1 accumulation at mitotic DNA breaks and the recruitment of TOPBP1 and CIP2A

During mitosis MDC1 recruits TOPBP1 and CIP2A to DNA breaks to tether them together throughout cell division, followed by break repair in the following G1-phase ^32,33^. To analyze the role of the PST repeat region in mitotic DNA break recognition and tethering, we genome edited the endogenous HaloTagged *MDC1* locus in U2OS cells to express MDC1 lacking the PST repeat region (Fig. 4A, Fig. S3A-C, Clones 33 and 37 are referred to as MDC1 ΔPST C1 and C2 throughout the manuscript). Cells expressing Halo-MDC1 ΔPST grew at a similar rate as their parental cells indicating that the deletion of the PST repeat region of MDC1 did not result in a proliferation defect (Fig. S3D). In mitotic control cells expressing wildtype MDC1, TOPBP1 was robustly recruited to DNA breaks marked by γH2AX (Fig. 3B). In contrast, both MDC1 and TOPBP1 failed to accumulate at DNA breaks in cells expressing MDC1 lacking the PST repeat region (Fig. 3B). Importantly, the formation of mitotic γH2AX foci was unaffected in cells expressing MDC1 ΔPST (Fig. 3B). TOPBP1 accumulated in large foci in mitotic cells expressing MDC1 ΔPST, but they did not overlap with γH2AX signals and were also present in control cells not treated with zeocin consistent with previous observations (Fig. 3B), suggesting that these TOPBP1 foci are not related to DNA break repair mediated by γH2AX ^32^. Similarly, we found that the presence of the PST repeat region of MDC1 was required for the recruitment of CIP2A to DNA lesions in mitosis (Fig. 4C). However, MDC1 containing non-phosphorylatable (T>A) and phospho-mimicking (T>D) variants of the PST repeat region co-localized with CIP2A in mitotic cells, indicating that the phosphorylation of the PST repeat region does not regulate the recruitment of CIP2A to sites of DNA damage in mitosis. To further analyze the role of the PST repeat region in recruiting MDC1 to DNA lesions in mitosis, we carried out laser micro-irradiation (LMI) experiments in cells arrested in M-phase using nocodazole. Wildtype Halo-MDC1 was readily recruited to LMI induced DNA lesions in mitotic cells (Fig. 4D, Movie S19). In contrast, only small amounts of Halo-MDC1 ΔPST accumulated at these sites of DNA damage (Fig. 4D, Movie S20-21). In total, these observations demonstrate that the PST repeat region of MDC1 supports the accumulation of MDC1 at and the recruitment of TOPBP1 and CIP2A to DNA breaks in mitosis.

**Figure 4.**
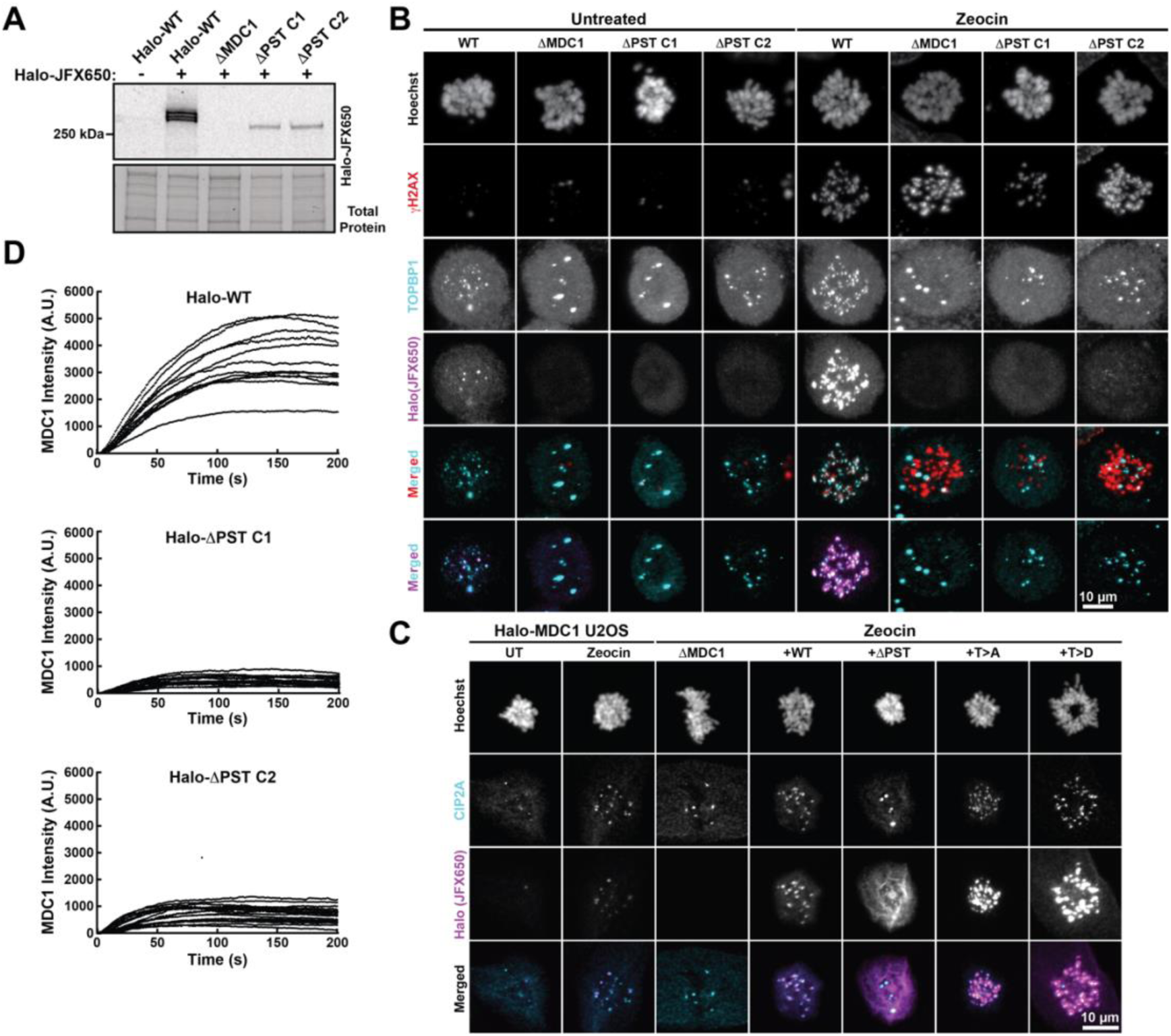
The PST repeat region of MDC1 is required for its accumulation at DNA breaks in mitosis and for the recruitment of TOBP1 and CIP2A. **(A)** Fluorescence gels of cell lysates from U2OS expressing HaloTagged MDC1 or MDC ΔPST from its endogenous locus labeled with JFX650 HaloTag-ligand. **(B)** Images of fixed, mitotic U2OS cells expressing HaloTagged MDC1 or MDC ΔPST from its endogenous locus untreated or treated with zeocin to introduce DNA breaks. Cells were labeled with JFX650 and Hoechst and immuno-stained with antibodies targeting TOPBP1 and γH2AX. **(C)** Images of fixed, mitotic U2OS cells expressing Halo-MDC1 and U2OS ΔMDC1 cells stably expressing HaloTagged MDC1 (JFX650) variants untreated or treated with zeocin. Cells were immuno-stained an antibody targeting CIP2A and stained with Hoechst to label DNA. **(D)** Quantification of the accumulation of knock-in HaloTagged MDC1 and MDC ΔPST at laser micro-irraditation induced DNA damage in living mitotic U2OS cells.

### Enrichment of MDC1 on mitotic chromatin leads to chromosome missegregation

We have identified a molecular switch that displaces MDC1 from mitotic chromatin via phosphorylation of the PST repeat region by CDK1-Cyclin B. Because the PST repeat region is able to form multivalent interactions with chromatin, we hypothesized that MDC1 has to be displaced from mitotic chromatin to prevent non-specific crosslinking of condensed chromosomes, since any attachments outside of those mediated by centromeric cohesion, like telomere fusions, can lead to chromosome missegregation. To test this hypothesis, we carried out timelapse imaging of mitotic progression in MDC1 knock-out cells stably expressing different HaloTagged MDC1 variants. In cells expressing wildtype MDC1 or MDC1 ΔPST the median time from nuclear envelope breakdown (NEB) to anaphase onset was ∼35 minutes (Fig. 5A,B, Movies S22-23). In contrast, expression of MDC1 with non-phosphorylatable (T>A) and phospho-mimicking (T>D) TP sites in the PST repeat region resulted in an increased time cells remained in mitosis, suggesting delays in chromosome alignment sufficient to satisfy the spindle assembly checkpoint (Fig. 5A,B, Movies S24-28). It is important to note that in these experiments wildtype MDC1 and MDC1 T>D accumulated on chromatin in mitotic cells (Fig. 5A), in contrast to earlier observations where wildtpe MDC1 and MDC1 T>D did not associate with mitotic chromatin (Fig. 1L). It is likely that MDC1 is expressed at higher levels from the doxycycline inducible cassette, overcoming the reduced affinity of phosphorylated wildtype MDC1 and MDC1 T>D for chromatin. Consistent with the hypothesis that T>A and T>D MDC1 interfere with chromosome alignment, the formation of a metaphase plate with completely aligned chromosomes was often delayed in cells expressing T>A and T>D variants of MDC1 (Fig. 5A, Movies S24-28). We also observed chromosome missegregation events ranging from the formation of chromosome bridges (Fig. 5A, Movie S25,28) to a complete failure of separating the chromosome masses during anaphase (Fig. 5A, Movie S26,27). The frequency of chromosome missegregation events was the highest for the T>A variant (∼46%), followed by the T>D allele (∼20%), and wildtype MDC1 (∼13%) (Fig. 5C), suggesting that chromosome missegregation is correlated with the affinity of the PST repeat region for chromatin. Consistent with this interpretation, chromosome missegregation events were rarely observed in cells expressing MDC1 ΔPST, which is completely excluded from mitotic chromosomes (Fig. 5A,C). In addition, the frequency of chromosome missegregation events was positively correlated to the expression level of MDC1 (Fig. 5C). In total, these observations demonstrate that accumulation of MDC1 on mitotic chromosomes can lead to inappropriate chromosome attachments that result in chromosome missegregation. This suggests that MDC1 has to be displaced from mitotic chromatin to prevent chromosome missegregation triggered by multivalent chromatin interactions of its PST repeat region.

**Figure 5.**
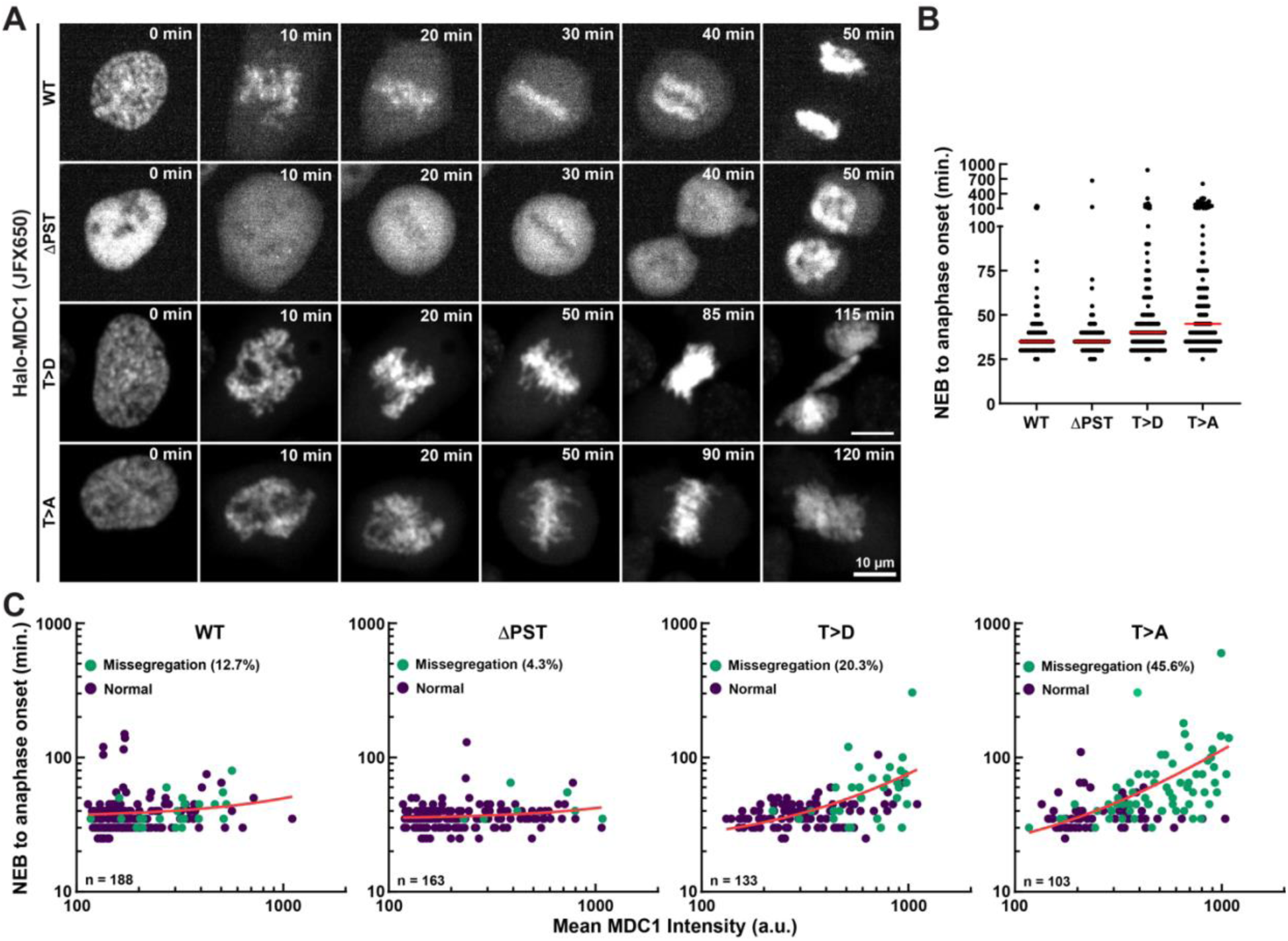
Accumulation of MDC1 on mitotic chromosomes leads to chromosome missegregation. **(A)** Still images from live cell timelapse imaging of U2OS ΔMDC1 stably expressing HaloTagged MDC1 variants labeled with JFX650. **(B)** Quantification of the time from nuclear envelope breakdown to anaphase onset of U2OS ΔMDC1 cells stably expressing HaloTagged MDC1 variants. **(C)** Quantification of the time from nuclear envelope breakdown to anaphase onset, the chromosome segregation phenotype, and Halo-MDC1 expression level of U2OS ΔMDC1 cells stably expressing HaloTagged MDC1 variants labeled with JFX650. The chromosome segregation phenotype was determined by manual inspection.

### The PST repeat region of MDC1 is sufficient for chromatin tethering

Our results demonstrate that accumulation of MDC1 on mitotic chromosomes acts as a molecular glue that counteracts proper chromosome segregation (Fig. 5). This phenotype was most pronounced in cells expressing the non-phosphorylatable (T>A) variant of MDC1, which is expected to have the highest affinity for chromatin in mitotic cells. This suggests that the ability of MDC1 to cause chromosome missegregation scales with its affinity for chromatin (*i.e.* the strength of the glue). To test whether local enrichment of the non-phosphorylatable (T>A) PST repeat region of MDC1 was sufficient to result in non-centromeric chromosome attachments and chromosome missegregation, we fused the wildtype and T>A PST repeat region to the Lac inhibitor (LacI) and GFP and expressed these constructs from a doxycycline inducible expression cassette in U2OS cells carrying a Lac operator (LacO) array on chromosome 1 containing more than 50,000 LacI binding sites ^56^. This approach allowed us to locally enrich and simultaneously visualize the PST repeat constructs at a single chromosomal locus (Fig. 6A). We induced expression of the LacI-PST repeat fusions and imaged mitotic cells using time lapse microscopy to determine whether the PST repeat region was sufficient to trigger missegregation of the chromosome carrying the LacO array (Fig. 6A). In cells expressing the wildtype PST repeat region fused to LacI, sister chromatids carrying the LacO array were equally segregated to the daughter cells with a single signal being detected within each chromosome mass upon anaphase onset (Fig. 6B,C, Movie S29). In contrast, in the presence of the T>A LacI-PST repeat fusion, we frequently observed the LacO array signal in the cleavage furrow inducing failed cytokinesis (Fig. 6B,C, Movie S30,31), detected fragmented or lagging LacO arrays during anaphase (Fig. 6B,C, Movie S32), and observed unequal segregation of the LacO array (Fig. 6B,C, Movie S33). These observations demonstrate that the local enrichment of the T>A PST repeat region results in sister chromatid attachments that are sufficient to trigger chromosome missegregation.

**Figure 6.**
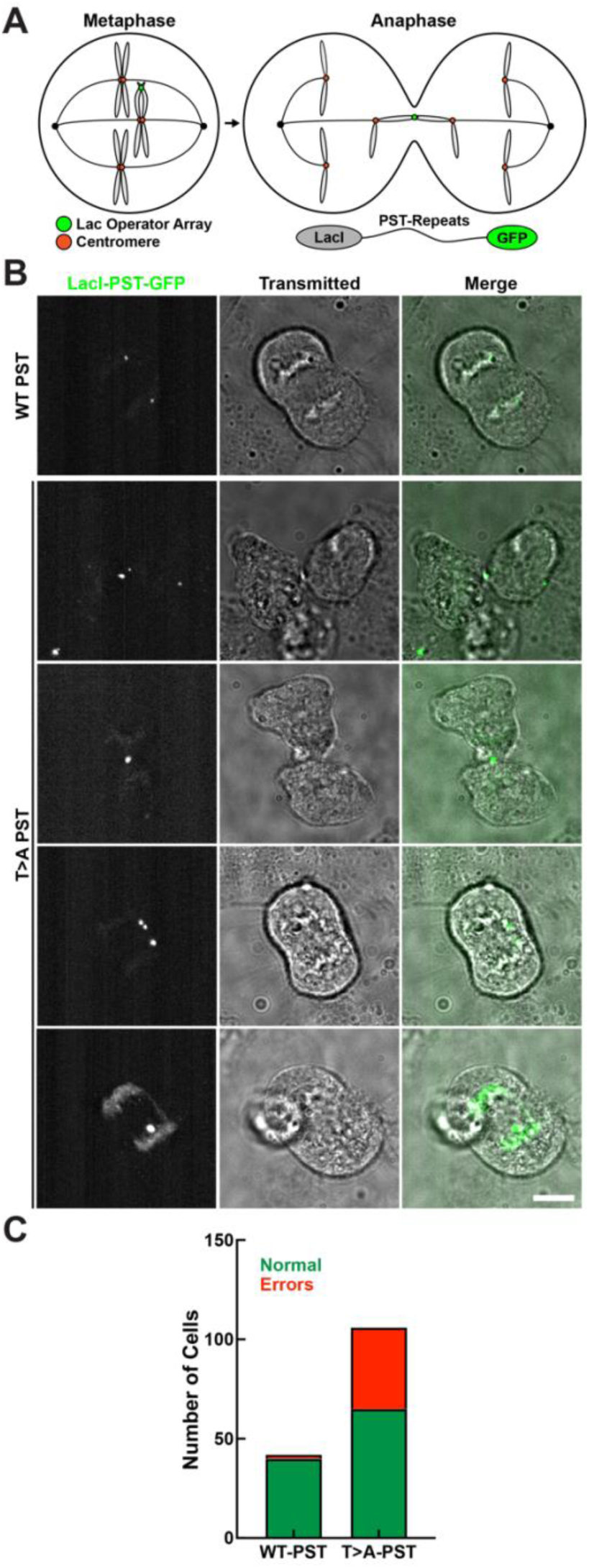
Local accumulation of the non-phosphorylatable T>A PST repeat region is sufficient to trigger chromosome missegregation. **(A)** Experimental design for local enrichment of the PST repeat region via the LacI-LacO interaction. **(B)** Still images from time lapse imaging of U2OS cells containing a single LacO array expressing wildtype and T>A LacI-PST repeat-GFP fusion proteins. **(C)** Quantification of the chromosome segregation phenotype of U2OS cells containing a single LacO array expressing wildtype and T>A LacI-PST repeat-GFP fusion proteins. Errors include lagging LacO arrays, LacO missegregation, and failed cytokinesis.

### The PST repeat region is required for homologous recombination

Previous observation by us and others have demonstrated that the PST repeat region of MDC1 contributes to DSB repair using the homologous recombination (HR) pathway. However, the molecular mechanism by which the PST repeat region contributes to HR remains unclear. To confirm that the PST repeat region is required for HR, we carried out clonogenic survival assays using the PARP inhibitor olaparib. Consistent with a defect in HR, cells expressing MDC1 ΔPST were highly sensitive to olaparib compared to control cells (Fig. 7A). To further dissect the contribution of the PST repeat region of MDC1 to HR, we analyzed RAD51 foci formation. While MDC1 ΔPST co-localized with zeocin induced DNA breaks marked by γH2AX, we observed greatly reduced formation of RAD51 foci in cells expressing MDC1 ΔPST (Fig. 7B). In contrast, RAD51 foci that co-localized with γH2AX and MDC1 were readily detected in control cells expressing wildtype Halo-MDC1 (Fig. 7B). Quantification revealed that deletion of either the PST repeat region or the BRCT domain of MDC1 almost completely eliminated RAD51 foci formation (Fig. 7C,D). In contrast, 53BP1 foci formation was largely unaffected by deletion of the PST repeat region of MDC1, while deletion of the BRCT domain eliminated 53BP1 foci formation (Fig. 7E,F). These observations demonstrate that the PST repeat region is not required for MDC1 localization to DNA breaks in interphase cells but is critical for a downstream step that eventually leads to RAD51 loading.

**Figure 7.**
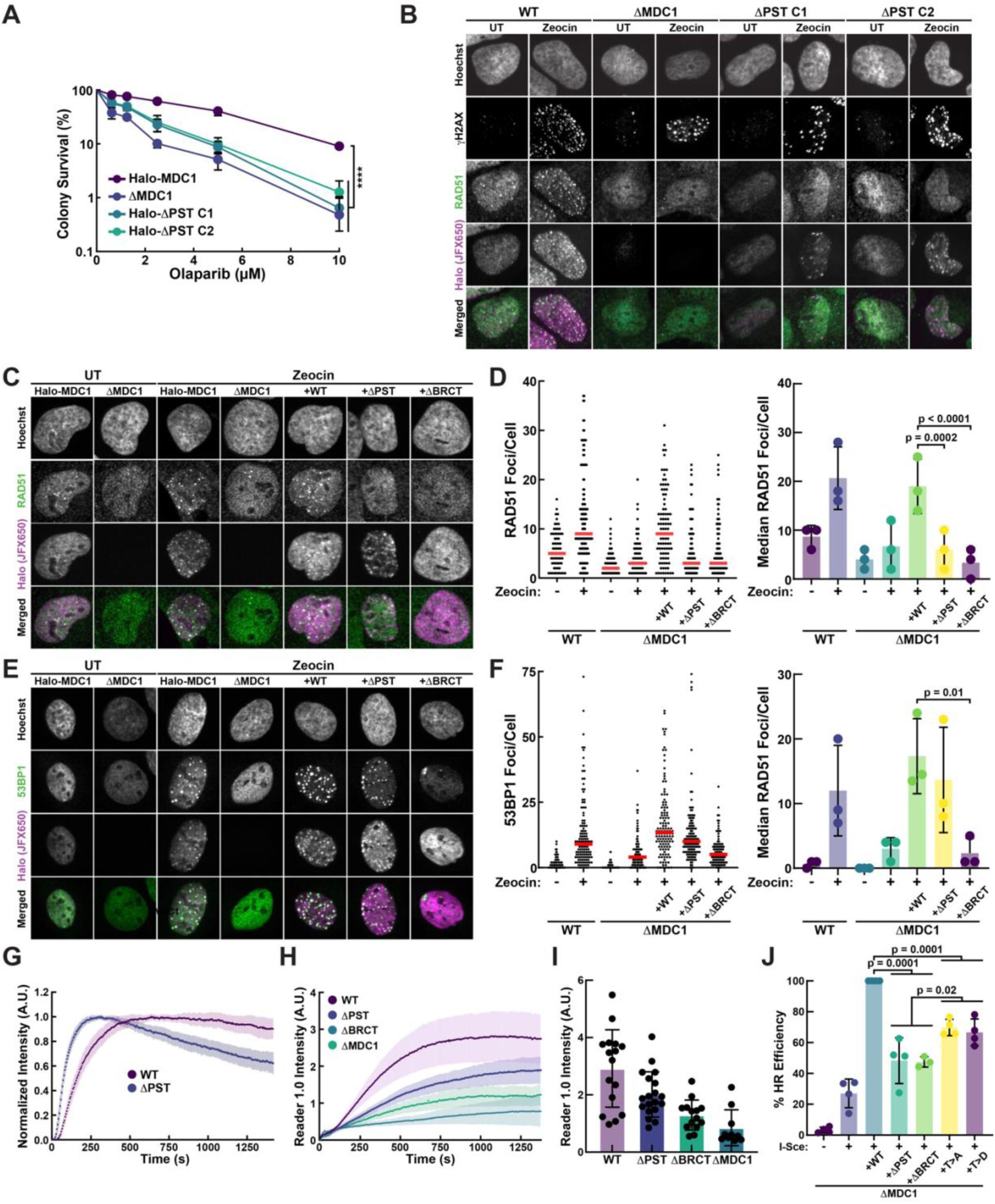
The PST repeat region is required for DNA break repair by homologous recombination. **(A)** Quantification of clonogenic survival assays using the PARP inhibitor olaparib of U2OS cells expressing HaloTagged MDC1 or MDC ΔPST from its endogenous locus. (N = 3 biological replicates plated in triplicate, Mean ± S.D., Two-way ANOVA) **(B)** Images of fixed interphase U2OS cells expressing HaloTagged MDC1 or MDC ΔPST from its endogenous locus untreated or treated with zeocin to induce DNA breaks. Cells were stained with JFX650 HaloTag-ligand and Hoechst, and immuno-labeled with antibodies targeting RAD51 and γH2AX. **(C)** Images of fixed interphase U2OS cells expressing Halo-MDC1 from its endogenous locus, MDC1 knock-out cells (ΔMDC1), and U2OS ΔMDC1 stably expressing HaloTagged MDC1 variants untreated or treated with zeocin to induce DNA breaks. Cells were stained with JFX650 HaloTag-ligand and Hoechst, and immuno-labeled with an antibody targeting RAD51. **(D)** Quantification of the number of RAD51 foci of the experiment in (C). An individual replicate is plotted on the left (red line = median, >100 cells per condition) and 3 biological replicates are plotted on the right (Median ± S.D., Two-way ANOVA). **(E)** Images of fixed interphase U2OS cells expressing Halo-MDC1 from its endogenous locus, MDC1 knock out cells (ΔMDC1), and U2OS ΔMDC1 cells stably expressing HaloTagged MDC1 variants untreated or treated with zeocin to induce DNA breaks. Cells were stained with JFX650 HaloTag-ligand and Hoechst, and immuno-labeled with an antibody targeting 53BP1. **(F)** Quantification of the number of 53BP1 foci of the experiment in (E). An individual replicate is plotted on the left (red line = median, >100 cells per condition) and 3 biological replicates are plotted on the right (Median ± S.D., Two-way ANOVA). **(G-H)** Quantification of the accumulation of **(G)** Halo-MDC1 and Halo-MDC1 ΔPST and **(H)** ubiquitin Reader1.0 at laser micro-irradiation induced DNA lesions in U2OS ΔMDC1 cells stably expressing indicated HaloTagged MDC1 variants (N = 10-19 cells, Mean ± 95% CI). **(H)** Quantification of the maximal accumulation of ubiquitin Reader1.0 at laser micro-irradiation induced DNA lesions in U2OS ΔMDC1 cells stably expressing indicated HaloTagged MDC1 variants (N = 10-19 cells, Mean ± 95% CI). **(G)** Quantification of HR efficiency using the DR-GFP reporter stably integrated into the AAVS1 locus of U2OS ΔMDC1 cells. I-Sce and MDC1 variants were transiently transfected (N = 3-4 biological replicates, Mean ± S.D., Two-way ANOVA).

Ubiquitin signaling mediated by RNF8 and RNF168 downstream of MDC1 is critical for the recruitment of HR (e.g. BRCA1) and NHEJ (e.g. 53BP1) effectors^19,57^. To analyze the role of the PST repeat region and BRCT domains of MDC1 in ubiquitin signaling we carried out experiments using the genetically encoded H2AK13/15-ubiquitin sensor Reader1.0 fused to eGFP and a nuclear localization sequence (NLS) to detect RN168 activity in living cells^58^. The NLS-Reader1.0-eGFP fusion protein was stably expressed using lentiviral transduction of MDC1 knock-out cells, and cell lines with inducible expression of HaloTagged wildtype MDC1, MDC1 ΔPST, and MDC1 ΔBRCT. To simultaneously analyze the recruitment of HaloTagged MDC1 and the Reader 1.0 to DNA lesions, we carried out laser micro-irradiation experiments to induce localized DNA damage. Consistent with ubiquitination being downstream of MDC1 recruitment, Reader 1.0 accumulation was delayed compared to MDC1 enrichment (Fig. 7G,H, Fig. S4A,B, Movie S34-37). We detected a small amount of chromatin ubiquitination in MDC1 knock out cells (Fig. 7G,H, Movie S37), suggesting that there must be a MDC1 independent chromatin ubiquitination pathway with limited activity. Compared to wildtype MDC1, ubiquitination detected by Reader1.0 was reduced to 70% and 44% in cells expressing MDC1 ΔPST or MDC1 ΔBRCT, respectively (Fig. 7I, Fig. S4C). This suggests that MDC1 lacking the PST repeat region, supports a reduced, yet substantial, amount of ubiquitin signaling.

Based on these observations, the inability of MDC1 ΔPST to support RAD51 loading could be due to reduced levels of ubiquitin signaling. However, ubiquitin signaling in cells expressing MDC1 ΔPST is sufficient for 53BP1 recruitment. Therefore, either the events necessary for RAD51 loading require a higher level of ubiquitination than 53BP1 recruitment, or another activity of the PST repeat region is required for HR progression. For instance, it is possible that multivalent interactions formed by the PST repeats with nucleosomes surrounding the DNA break are essential to tether DNA ends together throughout the prolonged HR process. To test whether modulating the affinity of the PST repeat region for chromatin alters HR efficiency, we carried out HR assays using the DR-GFP reporter stably integrated into the AAVS1 locus in MDC1 knock out cells transiently complemented with different MDC1 variants. In previous work we demonstrated that MDC1 ΔPST and MDC1 ΔBRCT had similar defects in HR using this approach ^45^. The MDC1 variants with non-phosphorylatable (T>A) and phospho-mimicking (T>D) mutations in the TP sites within the PST repeat region also had a significant defect in HR compared to wildtype MDC1 but supported slightly higher levels of HR compared to MDC1 ΔPST and MDC1 ΔBRCT (Fig. 7J). These observations suggest that altering the chromatin binding properties of the PST repeat region reduces the ability of MDC1 to facilitate homologous recombination, consistent with a role of chromatin tethering mediated by the PST repeat region in HR.

## DISCUSSION

The results described in this paper reveal in critical function of the PST repeat domain of MDC1 in DSB repair. We demonstrate that the PST repeats constitute a multivalent chromatin binding domain that in itself is sufficient to tether chromatin together. In concert with the BRCT domain of MDC1, which specifically recruits MDC1 to DNA breaks the chromatin tethering mediated by the PST repeats is required for DSB repair by homologous recombination. In mitotic cells, the affinity of the PST repeats for chromatin is downregulated by CDK1-Cyclin B mediated phosphorylation to prevent chromosome missegregation caused by non-specific adhesion of condensed mitotic chromosomes.

### The PST repeat region is a multivalent chromatin interaction domain with tunable affinity

Observations by Salguero *et al.* demonstrated the PST repeat domain of MDC1 interacts with the acidic patch of nucleosomes ^46^. Consistent with this model, our previous work using live cell single-molecule imaging demonstrated that MDC1 is constitutively associated with chromatin in human cells ^45^. This constitutive chromatin association depends on the PST repeat region of MDC1 but not the BRCT domain, which specifically binds γH2AX to target MDC1 to DNA breaks. In this work, we demonstrate that reducing the copy number of the PST repeats gradually decreases its association with chromatin. This suggests that each PST repeat contributes to a similar degree to chromatin binding and that the PST repeat region associates with multiple nucleosomes simultaneously. The ability to bind to multiple nucleosomes simultaneously allows the PST repeat region to tether together chromatin that is not connected by DNA, for instance in the context of a DNA break, or potentially to hold a sister chromatin in close proximity to facilitate HR.

Our work further demonstrates that in mitotic cells the affinity of the PST repeats for chromatin is regulated by CDK1-Cyclin B dependent phosphorylation of two conserved TP sites present in almost all of the individual 41 amino acid repeats. In undamaged cells this mitotic phosphorylation is sufficient to displace MDC1 expressed at endogenous levels from chromatin. However, overexpression of wildtype MDC1 or MDC1 with phospho-mimicking (T>D) mutations in the TP sites can result in chromatin association of MDC in mitotic cells, while deletion of the PST repeats results in complete exclusion of MDC1 from mitotic chromosomes. This demonstrates that phosphorylation of the TP sites in the PST repeats reduces the affinity of MDC1 for chromatin but does not completely eliminate binding. Consistent with this interpretation, the chromatin bound fraction of MDC1 with phospho-mimicking (T>D) mutations in interphase cells is significantly higher than that of MDC1 in which the entire PST repeat region has been deleted. While we did not observe phosphorylation of the TP sites within the PST repeats in interphase cells, it is interesting to consider the possibility that other posttranslational modifications of the repeats and chromatin context itself fine-tune the PST repeat-chromatin interaction in certain contexts (*e.g.* DNA replication) or predispose DSBs to repair by a specific pathway.

Importantly, we also address the functional relevance of mitotic phosphorylation of the PST repeat region of MDC1. The PST repeat region has the ability to form multivalent interactions with nucleosomes and therefore can crosslink chromatin, which could be detrimental in mitotic cells attempting to segregate condensed chromosomes. To facilitate accurate chromosome segregation, cells must eliminate all existing connections between sister chromatids or other chromosomes by removing cohesin complexes and alleviating chromosome entanglements using topoisomerases ^59^. Our experiments demonstrate that MDC1 with non-phosphorylatable TP sites in the PST repeat region (T>A) is strongly enriched on mitotic chromosomes and leads to a highly penetrant chromosome missegregation phenotype. In cells expressing MDC1 T>A we frequently observe complete failure to separate the chromosome masses during anaphase, consistent with non-specific adhesion of mitotic chromatin. This phenotype is correlated with the expression level of MDC1 and is also observed to a lesser degree in cells expressing wildtype MDC1 and MDC1 with phospho-mimicking mutation in the PST repeats. The severity of the phenotype is correlated with the affinity of the PST repeats for chromatin with very few missegregation events observed in cells expressing MDC1 ΔPST (4% of cells with missegregation), followed by wildtype MDC1 (expected to be fully phosphorylated, 13% of cells with missegregation), MDC1 T>D (20% of cells with missegregation), and MDC1 T>A, expression of which results in chromosome missegregation in almost half of the cells. In addition, our results demonstrate that enrichment of the non-phosphorylatable T>A variant of the PST repeat region at a single genomic locus (LacO array) is sufficient to trigger chromosome missegregation.

Consistent with a localized adhesion of sister chromatids containing the LacO array, we observed the LacO array signal lagging behind the main chromosome mass or stuck in the cleavage furrow when cells attempted to undergo cytokinesis (Fig. 5A). The observations that overexpressed WT or T>D MDC1 are tethered to mitotic chromatin and can also lead to chromosome missegregation suggests that MDC1 expression must be tightly controlled in mammalian cells. Protein levels need to be sufficiently high to facilitate DSB repair, yet low enough to limit deleterious events during chromosome segregation in mitosis which negatively impact genome instability.

In total our observations demonstrate that the PST repeat region of MDC1 is a multivalent chromatin interaction domain and its affinity for chromatin has to be downregulated in mitosis to prevent chromosome missegregation resulting from non-specific chromosome adhesion.

### MDC1 in mitotic DNA break tethering

In mitotic cells, MDC1 has been shown to contribute to tethering of DSBs to maintain their connectivity throughout cell division, allowing them to be repaired in the following G1-phase by 53BP1 dependent end-joining ^32^. To carry out this function, MDC1 recruits TOPBP1 to DNA breaks, via a direct interaction formed between a short motif in the N-terminus of MDC1 with the N-terminal region of TOPBP1 ^32^. TOPBP1 in turn recruits CIP2A, which is excluded from nuclei in interphase cells, to DNA breaks in mitosis ^33^. The MDC1-TOPBP1-CIP2A complex is not only critical to tether DNA breaks in mitotic cells but has also been shown to cluster together shattered chromosome fragments derived from micronuclei in mitotic cells, to assure their segregation into one of the daughter cells ^34,35^. It is important to note that conflicting results regarding the extent of the relationship between MDC1 and micronuclei clustering have been reported ^34,35^. Since both TOPBP1 and CIP2A are unequivocally required for clustering shattered chromatin fragments, and MDC1 is necessary for their recruitment to DNA breaks, we favor the model that MDC1 is essential for this process.

The molecular mechanism by which MDC1-TOPBP1-CIP2A facilitates DNA break tethering is unclear. One critical aspect is recruitment of the complex to DSBs mediated by the interaction of the MDC1 BRCT domain with γH2AX. One possibility is that either CIP2A, TOPBP1, or both form multimeric complexes that tether together MDC1 molecules bound to the two DNA ends. Based on our observations, we predict that tethering of chromatinized DNA fragments in mitosis is enhanced by the PST repeat region of MDC1. Importantly, our results demonstrate that the PST repeat region is required for accumulation of MDC1, TOPBP1, and CIP2A at DNA breaks in mitosis and future experiments uncoupling the break recruitment and chromatin tethering functions of the PST repeat region will be necessary to directly test this hypothesis. Our observations demonstrate that even phosphorylated wildtype MDC1 can tether mitotic chromatin when overexpressed. Therefore, it is likely that the PST repeats of MDC1 play an important role in chromatin tethering in context of mitotic MDC1-TOPBP1-CIP2A bound DNA breaks. If the PST repeat region of MDC1 physically tethers together DNA ends, CIP2A, which is a well-established regulator of protein phosphorylation via its interaction with protein phosphatase 2A and B56α ^60^, could be required to tune the affinity of the PST repeat region for chromatin by regulating its phosphorylation status to ensure effective DNA break tethering. CIP2A is typically thought of as an inhibitor of phosphatase activity and the precise role of CIP2A in regulating the phosphorylation of MDC1’s PST repeat region must be clarified in future work.

### Chromatin tethering by MDC1 is contributes to DNA break repair by homologous recombination

Previous work by others and our results have clearly demonstrated that the PST repeat region of MDC1 contributes to DSB repair by homologous recombination ^13,45,61^. However, the mechanism by which the PST repeats facilitate HR remains unclear. MDC1 contributes to both HR and NHEJ by initiating a ubiquitination cascade that results in the recruitment of BRCA1 and 53BP1, respectively ^11^. BRCA1 facilitates the recruitment of PALB2 and BRCA2 to DSBs, which loads RAD51 onto the exposed single-stranded DNA end ^63,65,67^. Our results demonstrate that ubiquitin signaling is reduced in cells expressing MDC1 lacking the PST repeat region. The signaling function of MDC1 ΔPST is sufficient for the recruitment of 53BP1 but not RAD51 loading. In addition, previous work by others has demonstrated that BRCA1 recruitment to DSBs is unaffected by deletion of the PST repeat region ^13^. This suggests that a molecular event downstream of BRCA1 recruitment but prior to RAD51 loading requires the PST repeat region of MDC1.

We propose that the PST repeat region of MDC1 tethers together the chromatin around DSBs to assure that the DNA ends and the sister chromatid are maintained in close proximity to facilitate break repair by HR. Consistent with this hypothesis, the HR efficiency in cells expressing phospho-mimicking and non-phosphorylatable variants of the PST repeat region is strongly reduced. This suggests that changing the affinity of the PST repeats for chromatin and thereby the tethering function of MDC1 impairs HR or that the affinity of the interaction must be dynamically controlled during this process. We hypothesize that when the PST repeat region is not present, DNA ends and the sister chromatid are not in close enough proximity to allow efficient RAD51 loading which is required for the formation of detectable RAD51 foci. Importantly both the T>A and T>D variants of MDC1 resulted in a reduction in HR efficiency, suggesting that dynamic regulation of chromatin binding by the PST repeat region contributes to chromatin tethering in the context of HR.

In total, the work presented here demonstrates that the multivalent and tunable chromatin binding activity of the PST repeat region of MDC1 contributes to DNA break repair by homologous recombination by tethering the chromatin surrounding DNA breaks in interphase cells and likely contributes to DNA break tethering in mitotic cells in concert with TOPBP1 and CIP2A.

## ACKNOWLEDGEMENTS

This work was supported by grants from the NIH (F32GM139292, K99GM149942) to J.R.H. and (DP2 GM142307) to J.C.S. We thank to Luke Lavis lab (HHMI Janelia Research Campus) for providing Janelia Fluor ligands. All flow cytometry data were obtained using instrumentation in the MSU Flow Cytometry Core Facility and we especially thank to Matthew Bernard and Daniel Vocelle for providing service regarding cell sorting and flow cytometry analysis. The flow cytometry facility is funded in part through the financial support of Michigan State University’s Office of Research & Innovation and Colleges of Osteopathic Medicine, Human Medicine, Veterinary Medicine, Natural Sciences, and Engineering. We thank to MSU IQ microscopy core facility for supporting our work.

## CONTRIBUTIONS

Conceptualization: J.R.H and J.C.S.; Experiments: J.R.H, M.M., and G.I.P.; Data Analysis: J.R.H., D.G.B., and J.C.S.; Writing: Original Draft: J.R.H. and J.C.S.; Writing: Review and Editing: J.R.H. and J.C.S.

## DATA AVAILABILITY

All primary data will be made available upon reasonable request by J.C.S.

## MATERIALS AND METHODS

### Cell lines and cell culture

U2OS, RPE1 p53+/+, and HEK293T cells were obtained from ATCC and cultured in RPMI (U2OS and RPE1) or DMEM (HEK293T) containing 10% fetal bovine serum (FBS) and 100 units/mL penicillin and 100 µg/mL streptomycin in a humidified incubator maintained at 37°C with 5% CO_2_. 3xFLAG-Halo-MDC1 knock-in and ΔMDC1 knockout U2OS cells were previously describedin Heyza *et al.*^45^. DR-GFP knock-in into the *AAVS1* locus of ΔMDC1 U2OS cells was previously described in Heyza *et al.*^45^. U2OS 2-6-3 cells containing the LacO array were a kind gift from David Spector and were maintained in medium containing 100 μg/mL hygromycin B^56^.

### Molecular cloning and plasmids

DNA oligonucleotides were purchased from IDT for cloning gRNA sequences. After annealing gRNAs were cloned into BpiI-digested pX330-U6-Chimeric_BB-CBh-hSpCas9 (a gift from Feng Zhang (Addgene plasmid # 42230))^62^. The gRNA/Cas9 and homology-directed repair donor plasmids used to introduce a 3xFLAG-Puro-HaloTag into the endogenous *MDC1* locus in RPE1 cells were previously described in Heyza *et al.*^45^. Sequences and cloning of Halo-MDC1 WT, ΔPST and ΔBRCT cloned into pRK2 (modified from pHTN HaloTag CMV-neo; Promega; #G7721) by Gibson assembly were previously described in Heyza *et al.*^45^. MDC1 T>D and T>A mutants were cloned into pRK2 using N- and C-terminal fragments flanking the PST repeat domain that were PCR amplified from WT MDC1 and two codon-optimized gene fragments encoding the PST domain for each mutant which were purchased as gBlocks from IDT. To generate PST truncation mutants, we first cloned into pRK2 a WT version of MDC1 containing a codon-optimized PST repeat region using the same approach used to generate the T>D and T>A mutants. Next, we PCR amplified the N-terminus of MDC1 and generated repeat truncations by amplifying the C-terminus using staggered forward primers in PST domain. N- and C-terminal fragments were cloned by Gibson assembly into pRK2 after linearization by inverse PCR. For tetracycline-inducible expression of MDC1 mutants, MDC1 cDNAs were amplified from pRK2 and cloned by Gibson assembly into pSBTet-Bla after linearization with SfiI (a gift from Eric Kowarz (Addgene plasmid # 60510)^55^. The Sleeping Beauty Transposase expression vector, SB100x, was a gift from Mark Groudine (Addgene plasmid # 127909)^64^. To knock-in the ΔPST mutant into the MDC1 locus, we generated a homology directed repair (HDR) donor plasmid by Gibson assembly containing left and right homology arms (One ordered as a gBlock and the other PCR amplified from MDC1 cDNA), C-terminal MDC1 cDNA (PCR amplified), and a puromycin resistance cassette (PCR amplified). The H2A K13/K15 ubiquitin sensor expression plasmid, NLS-Reader1.0-eGFP, was a kind gift from Robert Cohen and Tingting Yao^58^. For lentiviral expression, the NLS-Reader1.0-eGFP was PCR amplified and cloned by Gibson assembly into AgeI- and EcoR1-linearized FUGW (a gift from David Baltimore (Addgene plasmid # 14883)^66^. To knock-in inducible WT and T>A GFP-PST-LacI fusion proteins into the AAVS1 locus in U2OS 2-6-3 cells, we PCR amplified GFP and LacI from a GFP-LacI expression plasmid (a kind gift from Iain Cheeseman)^68^and codon-optimized WT and T>A PST repeat sequences and cloned them by Gibson assembly into MluI and SalI-linearized AAVS1-TRE3G-EGFP (a gift from Su-Chun Zhang (Addgene plasmid # 52343))^69^. The AAVS1-targeting gRNA plasmid, PX458-AAVS1, was a gift from Adam Karpf (Addgene plasmid # 113194). To express and purify GST-tagged PST repeats from E. coli, two gBlocks for each protein fragment were codon-optimized for expression in E. coli and were cloned by Gibson assembly into BamHI-linearized pGSTag (a gift from David Ron, (Addgene plasmid # 21877)^70^. All newly generated plasmids were validated by Sanger Sequencing or Next Generation Sequencing (Quintara Biosciences and Genewiz).

### Lentivirus

Lentiviral particles to express NLS-Reader1.0-eGFP were produced in HEK293T cells. HEK293T cells were seeded onto 10 cm plates until they reached ∼70% confluency. Cells were co-transfected with FUGW (NLS-Reader1.0-eGFP), pRSV-Rev, pMDLg/pRRE, and pMD2.G with FuGene6 at a 3:1 ratio of FuGene6:DNA in antibiotic-free medium and incubated overnight. The following day, transfection medium was removed and 10 mL of antibiotic-free medium was added. Virus was collected at 48- and 72-hours post-transfection, filtered through a 0.4 µm filter, and 1 mL aliquots frozen at −80°C. For lentiviral transduction, cells were seeded onto 10 cm plates and allowed to attach overnight. The following day, viral aliquots were warmed to 37°C, 8 µg/mL polybrene was added, and virus was added dropwise to each plate. Experiments were carried out with the pooled population of transduced cells.

### Genome-editing

To knock-in the ΔPST mutant in the endogenous HaloTagged *MDC1* locus, GFP-PST-LacI into the *AAVS1* locus, and 3xFLAG-Puro-HaloTag into the endogenous *MDC1* locus in RPE1 cells, Halo-MDC1 U2OS, U2OS 2-6-3, and RPE1 cells were transfected with 1 mg each of pX330 and the HDR donor overnight using FuGene6 at a 3:1 ratio of Fugene6:DNA in antibiotic-free medium. The following day, transfection medium was replaced with complete medium. Knock-in cells were selected for with 1 µg/mL (PST) or 1.5 µg/mL puromycin. For WT and T>A GFP-PST-LacI, a single knock-in clone survived selection. ΔPST knock-in clones were isolated in 96-well plates by single-cell dilution, screened using genomic PCR, and validated by in-gel fluorescent protein detection and Sanger sequencing. RPE1 Halo-MDC1 cells were transfected with a plasmid encoding GFP-Cre to recombine out the puromycin resistance cassette oriented between the 3xFLAG and HaloTag and sorted by fluorescence activated cell sorting into 96-well plates. Clones were screened by SDS-PAGE and genomic PCR.

### Phospho-PST antibody production

Sera containing phospho-specific antibodies were generated against the following peptides (Ac-DQPV(pT)PEPTSC-amide, Ac-CNRSSVK(pT)PE-amide) by Labcorp using two rabbits per target and a 118-day protocol. All bleeds from both rabbits were pooled and purified. Peptides containing the phosphorylated and unphosphorylated peptides were crosslinked to SulfoLink^TM^ colums (Thermo Scientific, 44999). Sera were first flowed over columns with the unphosphorylated peptide to deplete antibodies that recognized parts of the peptide not specific to the phosphorylated threonine. The flowthroughs from these columns were flown over the columns with crosslinked phosphorylated peptide, washed, and eluted using acidic and basic conditions. Fractions containing high protein concentrations from both elution conditions were pooled and dialyzed into PBS containing 50% glycerol, spun down at 20,000xg for 15 minutes, and supernatants were aliquoted, snap frozen in liquid nitrogen, and stored at −80°C.

## SDS-PAGE

Cells grown in 24-well or 6-well plates were labeled with 100 nM Halo-JF646 or JFX650 for 10 minutes at 37°C and unbound ligand removed by washing the samples with complete medium. Prior to lysis samples were washed once with PBS then lysed in each well with 2x Laemmli buffer (BioRad) with β-mercaptoethanol. After boiling samples at 95°C for 5 minutes, equal amounts of lysate were loaded onto 4-15% or 4-20% TGX stain-free polyacrylamide gels. Gels were run at 180V for ∼1 hour. In-gel fluorescent detection of Halo-MDC1 was performed using the Cy5.5 filter on a BioRad Chemidoc. Stain-free protein detection to monitor sample loading was done using a BioRad Chemidoc and the Stain-free filter after 45 second activation with UV.

### Colony survival assay

600-800 cells/well were plated in 6-well plates in complete medium and allowed to attach overnight. The following day, medium was removed and fresh medium containing increasing concentrations of olaparib (Selleckchem; S1060) was added for the duration of the experiment. Approximately 6-7 days after treatment, medium was removed, wells washed once with phosphate-buffered saline (PBS), and cells fixed/stained with crystal violet solution (20% ethanol, 1% w/v crystal violet). After removing staining solution, plates were washed with water and allowed to air-dry. Plates were imaged on a BioRad Chemidoc and colonies counted using OpenCFU^71^. Experiments were performed three times in triplicate.

### Timelapse imaging of MDC1 in mitosis

Timelapse imaging was carried out on a Yokogawa CellVoyager CQ1 Benchtop High-Content Analysis System using a 40x objective. Tet-inducible Halo-MDC1 cells were seeded onto 24-well glass bottom plates two days prior to imaging and MDC1 expression was induced with 2 μg/mL doxycycline. Cells were maintained at 37°C with 5% CO_2_ during the course of imaging. Multiple fields of view were randomly selected for each sample and 3D images were acquired every 5 minutes for ∼20 hours. Z-projected images were analyzed in ImageJ/Fiji where the mean MDC1 fluorescence intensity for each analyzed cell was measured prior to mitotic entry and time to anaphase onset was measured by manually counting the number of frames starting with the first frame where dissolution of the nuclear envelope was observed until chromosome segregation or attempted segregation was initiated. At least 100 cells were analyzed for each mutant.

### Microscopy

Experiments were carried out on two microscope systems. The first system was a Olympus IX83 inverted microscope equipped with an environmental control unit, a cellTIRF illuminator (405, 488, 561, and 641 nm laser lines), an Excelitas X-Cite TURBO LED light source, Olympus UAPO 100x (1.49 NA) and 60x (1.5 NA) TIRF objectives, a cellFRAP with a 100 mW 405 nm laser, and 2 Andor iXon 897 Ultra EMCCD, Hamamatsu Orca-Fusion BT sCMOS, or Hamamatsu Orca-Quest qCMOS cameras, and operated using Olympus cellSense software. The second system was a 3i spinning-disc confocal (Yokogawa) microscope with SoRa modality equipped with incubation chamber (temperature, humidity and CO_2_ controlled), four laser lines (445, 488, 561 and 638 nm), Zeiss C PlanApo 63x/1.42 NA objective and Hamamatsu Orca-Fusion BT sCMOS or Hamamatsu Orca-Quest qCMOS camera.

### Live-cell imaging

All live-cell imaging experiments were carried out at 37°C, 5% CO_2_ and 95% humidity-controlled environment to mimic the conditions used for cell culturing. Cells were seeded into 24-well glass bottom plates for live-cell and live-cell single-molecule imaging. For imaging of densely labeled Halo-MDC1 mutants in mitosis, ΔMDC1 cells were transfected with plasmids encoding Halo-WT, T>A, or T>D MDC1 in 6 well plates two days prior to seeding for imaging using FuGene6. 60,000 cells were seeded for imaging and allowed to attach overnight. The following day, 100 ng/mL nocodazole was added to each well and cells were incubated for ∼16 hours. Immediately prior to imaging, Halo-MDC1 and DNA was labeled with 250 nM Halo-JF646 and Hoechst for ∼10 minutes in the presence of nocodazole, washed three times with complete medium, and fresh medium containing nocodazole was added. For live-cell single-molecule imaging of Halo-MDC1 mutants in interphase, cells were transfected using FuGene6 with plasmids encoding each mutant in 6-well plates and subsequently seeded for imaging or were nucleofected (PST truncation mutants) using a Lonza 4D Nucleofector and plates onto 24-well glass bottom plates. For single-molecule imaging in mitosis, samples were plated onto 24-well glass bottom plates and allowed to attach overnight. ∼24 hours later, 100 ng/mL nocodazole was added to each well and cells were incubated for ∼16-20 hours prior to imaging. Immediately prior to imaging, Halo-MDC1 was labeled with either Halo-JF646 or Halo-JFX650 (0.5 – 4 nM for 1 minute). Excess ligand was removed by washing the samples three times with complete medium, incubated for 10 minutes at 37 °C, and fresh complete medium added. Imaging was done using either the 100x or 60x objective and the 640 nm laser line with highly inclined laminated optical sheet (HILO) illumination. Images were acquired at 138 - 198 fps for at least 1000 frames.

### Single-molecule imaging analysis

Live-cell single-molecule imaging movies were analyzed using single-particle tracking (SPT) in MATLAB 2019a with a version of SLIMfast that allows for analysis of TIFF files ^49^. SPT setting use for analysis: Exposure Time = 5 – 7.20 ms, NA = 1.49 or 1.5, Pixel Size = 0.16 mm (Andor iXon Ultra) or 0.1083 mm (Hamamtsu ORCA Fusion BT), Emission Wavelength = 664 nm, *D*_max_ = 5 mm^2^/s, Number of gaps allowed = 2, Localization Error = −5, Deflation Loops = 0. Next, particle tracks were analyzed using SpotOn in MATLAB to obtain diffusion coefficients and the fraction of bound and freely diffusing particles. SpotOn settings used for the analysis of SPT files are as follows: TimeGap = 5 – 7.20 ms. dZ = 0.700 mm, GapsAllowed = 2, TimePoints = 8, JumpsToConsider = 4, BinWidth = 0.01 mm, PDF-fitting, D_Free_2State = [0.5 25], D_Bound_2State = [0.0001 0.5]. At least 20 cells for each of three biological replicates were imaged for all live-cell single molecule imaging experiments. Summary data for each biological replicate were plotted in GraphPad Prism.

### Laser microirradiation

Laser microirradiation (LMI) experiments were carried out using the Olympus ix83 microscope described above. Cells were seeded onto 24-well glass bottom plates for LMI experiments. For tet-inducible MDC1 mutants, 2 μg/mL doxycycline was added 1-2 days prior to imaging to induce protein expression. For dual-color imaging of Halo-MDC1 and NLS-Reader1.0-eGFP, cells were transduced with NLS-Reader1.0-eGFP lentiviral particles in 10 cm plates, were subsequently seeded onto 24-well glass bottom plates, and Halo-MDC1 expression induced with doxycycline. For LMI carried out on mitotic cells, mitotic cells were enriched by adding 100 ng/mL nocodazole to each well for ∼16-20 hours prior to imaging. On the day of imaging, Halo-MDC1 was labeled with 100-150 nM Halo-JFX650 and simultaneously pre-sensitized with Hoechst (1 μg/mL) for ten minutes (in the presence of nocodazole for mitotic samples). After labeling, cells were washed three times with complete medium, incubated for 10 minutes at 37°C, followed by the addition of fresh complete medium. Cells were maintained at 37°C with 5% CO_2_ during the course of imaging. Cells were irradiated using an interpolated line at 20% laser power with a 20 ms pulse. Images were acquired using a 60x objective and fluorescence imaged after excitement with the 475 or 630 nm LED light source at various rates (*e.g*., every 0.5 s, 1 s, 5 s). Images were converted to TIFF files and Halo-MDC1 and NLS-Reader1.0-GFP recruitment were quantified in Fiji by placing an ROI around the irradiated line and measuring mean intensity over time. From these data, the mean fluorescence at baseline was subtracted to calculate the relative increase in fluorescence over time. Normalized fluorescence intensity values were determined by setting the highest intensity frame for each cell to 1. Similarly, for normalizing the intensity values for the average of all cells combined for each group, the frame with the highest mean intensity was set to one. Data were plotted in GraphPad Prism.

### Purification of GST-PST repeats and in vitro kinase assay

Plasmids encoding WT and T>A GST-PST repeats were transformed into OneShot^TM^ BL21(DE3) cells (Invitrogen). Bacterial cultures were grown at 37°C at 220 rpm to an OD600 of 0.7. Protein expression was induced with 1 mM IPTG and cultures grown for ∼16 hours at 18 °C with shaking at 180 rpm. Cells were harvested by centrifugation at 5000xg for 5 minutes at 4°C and frozen at −80°C. Pellets were resuspended in wash and lysis buffer (250 mM NaCl, 1 mM DTT (added fresh) in PBS). Lysozyme was added at a concentration of 0.5 mg/mL and samples were sonicated (90 seconds, 35% amplitude, 10 second pulses, 20 second pause, Fisherbrand Model 505, 0.5 inch tip) in an ice water bath. Lysates were cleared by centrifugation at 40,000xg for 30 minutes at 4°C. 2 mL of Glutathione Sepharose (GE Healthcare; 17-5132-01) was equilibrated by washing three times with 20 mL of wash and lysis buffer and spun down at 1000 x g for two minutes between washes. After clearing, lysate was added to the GST agarose and rotated for one hour at 4°C. Samples were spun down at 1000xg for two minutes, lysate removed, and samples washed 3x with 10 mL of wash and lysis buffer. After washing, 4 mL of elution buffer (50 mM Tris pH8.1, 75 mM KCl, 10 mM Glutathione (added fresh)) was added to each sample, rotated for 10 minutes at 4°C, and eluted protein collected. GST-PST repeats were additionally purified using size exclusion chromatography using a Superdex 200 column into 50 mM Tris pH 7.0, 150 mM KCl, 1 mM DTT. Fractions were run on an SDS-PAGE gel and peak fractions combined and concentrated. Concentrated protein was supplemented with 50% glycerol, frozen in liquid nitrogen and stored at −80°C. Protein concentration was measured by absorption spectroscopy using ɛ280nm = 46,090 M-1 cm-1 which was calculated using the GST-PST sequence and ExPASy ProtParam.

*In vitro* kinase assays with purified Cyclin B1/CDK1 and WT/T>A GST-PST repeats were performed following a protocol described in Wang *et al.*^72^. Purified human Cyclin B/CDK1 was purchased from Invitrogen (PV3292). 2.5 μg GST-PST and ∼380 ng Cyclin B/CDK1 were incubated in 1x kinase buffer (0.25 M Tris-HCl, pH 7.5, 0.05 M MgCl_2_, 0.5 M EDTA, 10 mM DTT), supplemented with 100 μCi/mmol of [γ-^32^P] ATP (PerkinElmer) and incubated at 30°C for one hour. The reaction was stopped by adding 2x Laemmli buffer containing β-mercaptoethanol. Samples were incubated at 60°C for 10 minutesthen loaded onto SDS-PAGE gels. Samples were visualized using autoradiography with an Amersham Typhoon (GE).

### DR-GFP Assay

DR-GFP assays were performed three-four times and fluorescence detected by flow cytometry. ΔMDC1 DR-GFP knock-in cells were nucleofected using a Lonza 4D nucleofector in RPMI plus 50 mM HEPES with 400 ng I-SceI plasmid (a gift from Maria Jasin (Addgene plasmid # 26477)^73^ and 600 ng of plasmids encoding Halo-MDC1 mutants or 3xFLAG-Halo-NLS (previously described Klump *et al.*^74^). Cells were seeded into 6-well plates. 48 hours after nucleofection, HaloTag was labeled with 150 nM Halo-JFX650 and cells were trypsinized and collected. Samples were run on a BD Accuri C6 cytometer and at least 5,000 JFX650-positive events collected for each sample and analyzed in FCS Express 7.

### Immunofluorescence

For immunofluorescence (IF), cells were seeded onto glass coverslips in 6-well plates and allowed to attach for 1 day. For PST truncation mutants, ΔMDC1 cells were nucleofected with each Halo-MDC1 plasmid using a Lonza 4D nucleofector in serum- and antibiotic-free RPMI medium supplemented with 50 mM HEPES buffer solution (GIBCO; 15630-106) immediately prior to seeding. For tet-inducible MDC1, protein expression was induced by adding 2 μg/mL doxycycline for 2 days prior to proceeding with IF. For mitotic samples, cells were synchronized at G2/M with CDK1i (9 μM; Selleckchem; S7447) for ∼16-20 hours. HaloTagged protein was labeled at the time of release from CDK1i by adding 100 nM Halo-JFX650 and incubating for 10 minutes. Samples were washed three times to remove unbound ligand and released into medium containing S-Trityl-L-cysteine (STLC) (10 μM; Enzo Life Sciences, ALX-105–011-M500) and ± Zeocin (20 μg/mL; Invitrogen; R250011) for 1 hour. For images of different stages of mitosis, samples were released from CDK1i into complete medium ± Zeocin and fixed at 60, 90, or 120 minutes post-release. For interphase samples, HaloTagged protein was labeled with 100 nM Halo-JFX650 for 10 minutes followed by three washes with complete medium and subsequent incubated for 4 hours ± Zeocin (20 μg/mL). Cells were washed once with PBS and fixed with 4% formaldehyde or ice-cold 100% methanol (for TOPBP1 or p-PST staining) for 10 minutes at room temperature. Fixative was removed by washing samples once with PBS followed by permeabilization with 0.2% Triton-X 100 in PBS. Samples were washed twice with antibody dilution buffer (ABDIL) (3% BSA in PBS-Tween 20 (PBS-T)) and blocked in ABDIL for 30-60 minutes. Samples were incubated with primary antibodies for 1 – 1.5 hours at room temperature, washed three times with PBS-T, and incubated with secondary antibodies for 1 hour at room temperature. After washing three times with PBS-T, DNA was labeled with Hoechst, coverslips washed once with PBS, mounted onto glass slides with ProLong Diamond Antifade Mountant (Invitrogen; P36970) and sealed with nail polish. Samples were imaged on a 3i spinning-disc microscope with a 63x objective. Antibodies: anti-γH2AX (Millipore; JBW301; 05-636) (1:1,000); anti-TOPBP1 (Bethyl; A300-111A) (1:500); anti-CIP2A (Santa Cruz; sc-80659) (1:500); anti-53BP1 (Novus; NB100-304) (1:1000); anti-p-PST #1 (Stock concentration = 3 mg/mL) (1:3,000); anti-p-PST #2 (Stock concentration = 6.2 mg/mL) (1:6,000); Goat anti-mouse IgG Antibody, Cy3 conjugate (Millipore; AP124C) (1:500); Cy3 goat anti-rabbit IgG (H+L) (Invitrogen; A10520) (1:500); Goat anti-mouse IgG (H+L) AF488 (Invitrogen; A32723) (1:500 for CIP2A; 1:1,000 for γH2AX). Goat anti-rabbit IgG (H+L) (Invitrogen; A11034) (1:500).

### Imaging of LacI-PST-GFP fusions in U2OS 2-6-3 cells

U2OS 2-6-3 cells expressing inducible WT and T>A GFP-PST-LacI knocked into the *AAVS1* locus were seeded onto 24-well glass bottom plates and allowed to attach overnight. GFP-PST-LacI expression was induced with 1 μg/mL doxycycline. Timelapse images were acquired using a 63x objective on the 3i spinning disk described above. Cells were not synchronized and fields of view were selected based upon the presence of mitotic cells. Z-stack images for each XY position were acquired every 3 minutes for 1.5 - 2 hours. ≥ 10 mitotic cells for each mutant were imaged per replicate on three separate days. The cell division phenotypes were assessed by manually inspecting each cell undergoing anaphase and analyzing the segregation of the LacO array signal as well as the overall cell morphology after cell division. Failed cytokinesis was apparent by persistent connection bridging the two cell bodies or by opening of the cleavage furrow leading to a bi-nucleated cell.

## Supplemental Material

**Supplemental Figure 1.**
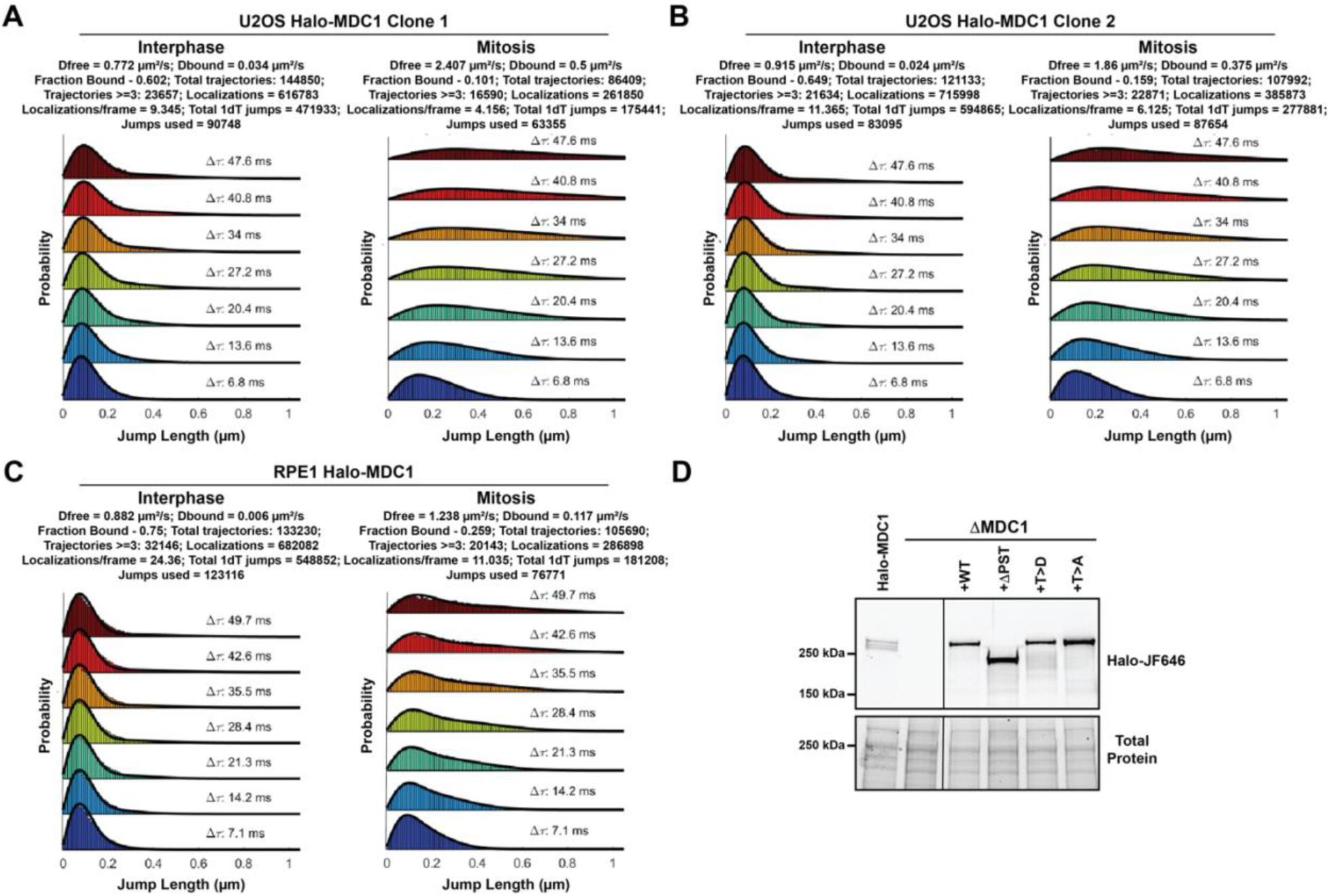
**(A-C)** Displacement histograms and two state fit of single particle tracking analysis of Halo-MDC1 expressed from its endogenous locus in **(A,B)** U2OS cells and **(C)** RPE1 cells. **(D)** Fluorescence gel of cell lysates from U2OS expressing Halo-MDC1 from its endogenous locus and U2OS ΔMDC1 cells transiently expressing the indicated HaloTagged MDC1 variants labeled with JF646 HaloTag-ligand.

**Supplemental Figure 2.**
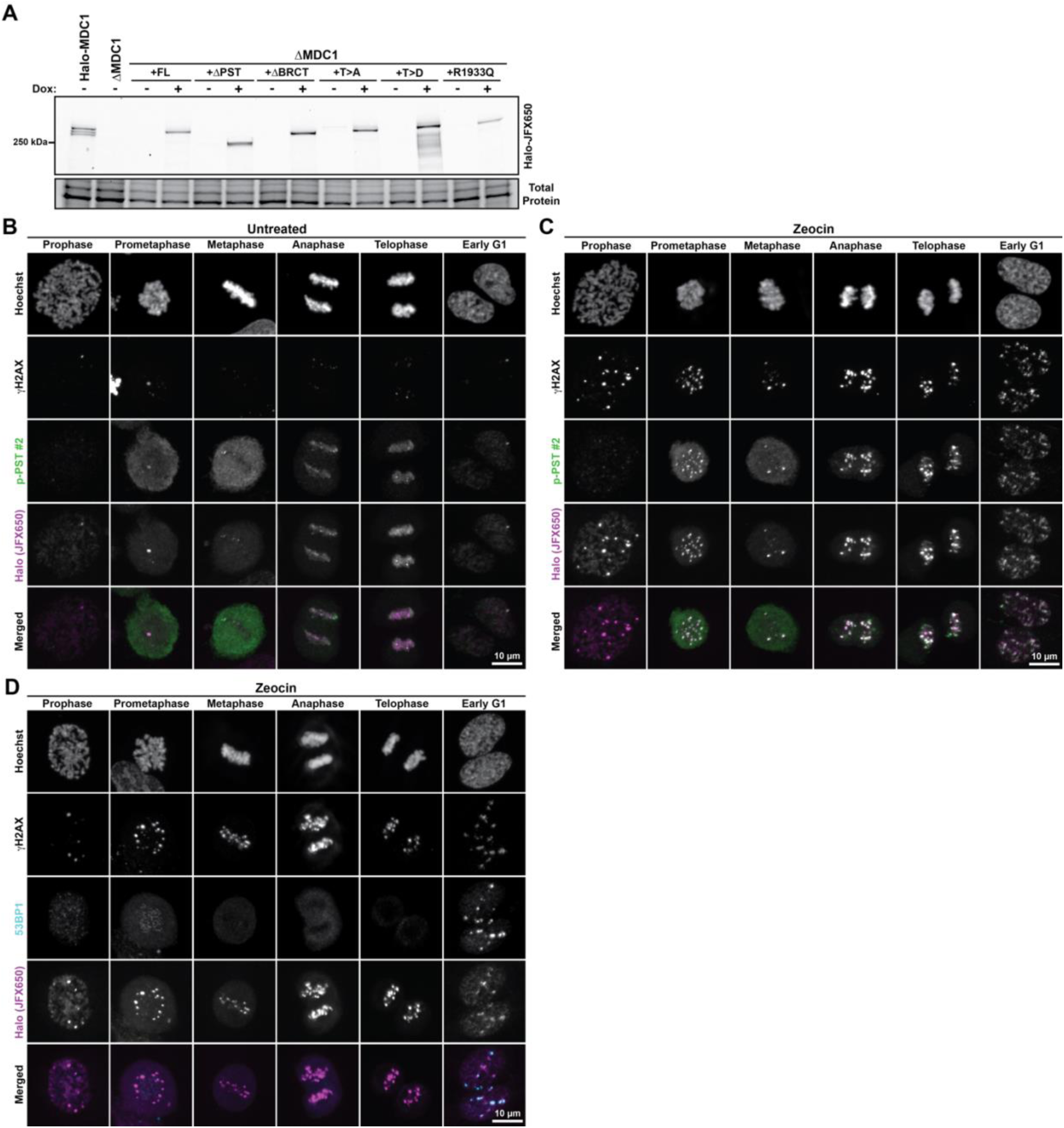
**(A)** Fluorescence gel of cell lysates from U2OS expressing Halo-MDC1 from its endogenous locus and U2OS ΔMDC1 cells expressing doxycycline (Dox.) inducible HaloTagged MDC1 variants from an expression cassette integrated using Sleeping Beauty transposase. The HaloTag was labeled with JFX650 HaloTag-ligand. **(B-C)** Images of fixed U2OS cells expressing Halo-MDC1 (JFX650) **(B)** untreated or **(C)** treated with zeocin to induce DNA breaks in various phases of mitosis and early G1-Phase. Cells were immuno-stained with p-PST antibody #2 and a γH2AX antibody and Hoechst to label DNA. **(D)** Images of fixed U2OS cells expressing Halo-MDC1 (JFX650) treated with zeocin to induce DNA breaks in various phases of mitosis and early G1-Phase. Cells were immuno-stained with antibodies against 53BP1 and γH2AX and Hoechst to label DNA.

**Supplemental Figure S3.**
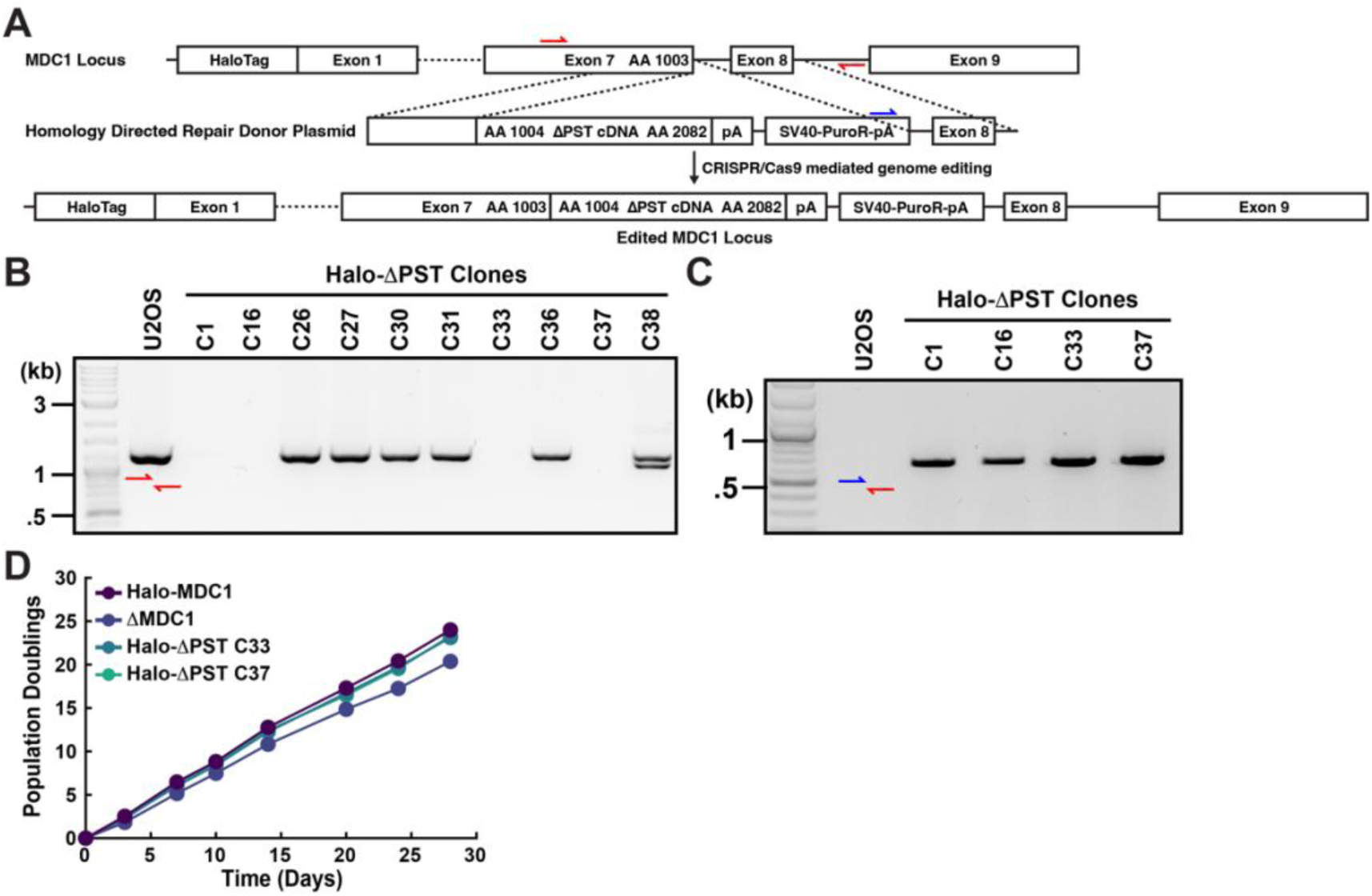
**(A)** Genome editing strategy to knock-in a cDNA encoding MDC1 lacking its PST repeat region into the endogenous *MDC1* locus. Colored arrows indicate primer positions. **(B-C)** PCR using genomic DNA from clonal U2OS cell lines subjected to genome editing using the indicated primer pairs. In **(B)** successful genome editing of all alleles is expected to lead no product amplification due to the large insertion of the cDNA and selection cassette. In **(C)** successful insertion of the donor sequence is validated using a reverse primer oriented downstream of the right homology arm and a forward primer in the donor. **(D)** Growth curve of U2OS cells expressing Halo-MDC1 and Halo-MDC1 ΔPST from the endogenous *MDC1* locus, and MDC1 knock-out cells (ΔMDC1).

**Supplemental Figure S4.**
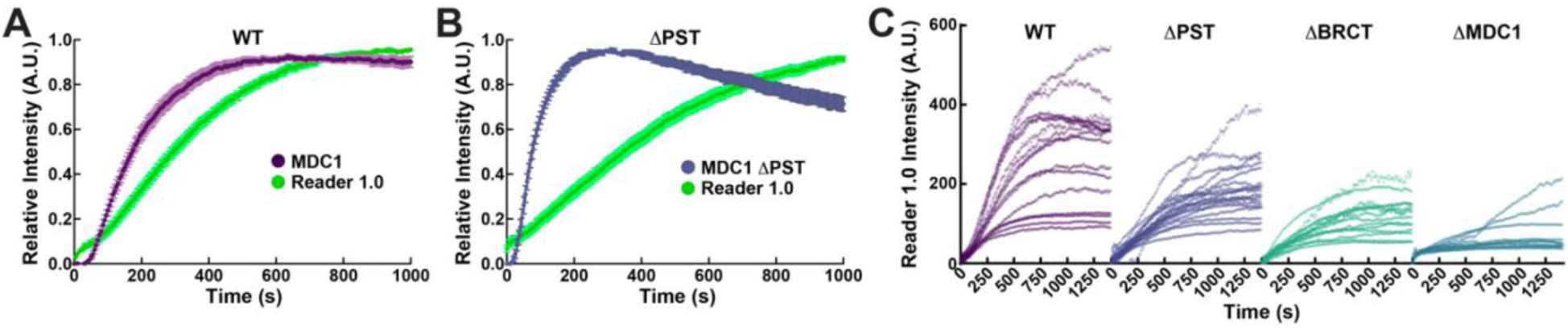
**(A-B)** Quantification of the accumulation of **(A)** Halo-MDC1 or **(B)** Halo-MDC1 ΔPST labeled with JFX650 HaloTag-ligand and ubiquitin Reader1.0 at laser micro-irradiation induced DNA lesions in U2OS ΔMDC1 cells stably expressing indicated HaloTagged MDC1 variants (N = 16-19 cells, Mean ± S.E.M.). **(C)** Accumulation of ubiquitin Reader1.0 at laser micro-irradiation induced DNA lesions in U2OS ΔMDC1 cells stably expressing indicated HaloTagged MDC1 variants. Each trace represents an individual cell.

## Supplemental Movie Legends

**Movie S1.** Live-cell single-molecule imaging movie of endogenous 3xFLAG-Halo-MDC1(Clone 1) U2OS cells in interphase. Images acquired at 138 frames per second.

**Movie S2.** Live-cell single-molecule imaging movie of endogenous 3xFLAG-Halo-MDC1(Clone 1) U2OS cells in nocodazole-arrested mitotic cells. Images acquired at 138 frames per second.

**Movie S3.** Live-cell single-molecule imaging movie of endogenous 3xFLAG-Halo-MDC1(Clone 2) U2OS cells in interphase. Images acquired at 138 frames per second.

**Movie S4.** Live-cell single-molecule imaging movie of endogenous 3xFLAG-Halo-MDC1(Clone 2) U2OS cells in nocodazole-arrested mitotic cells. Images acquired at 138 frames per second.

**Movie S5.** Live-cell single-molecule imaging movie of endogenous 3xFLAG-Halo-MDC1 RPE1 cells in interphase. Images acquired at 141 frames per second.

**Movie S6.** Live-cell single-molecule imaging movie of endogenous 3xFLAG-Halo-MDC1 RPE1 cells in nocodazole-arrested mitotic cells. Images acquired at 141 frames per second.

**Movie S7.** Live-cell single-molecule imaging movie of transiently expressed 3xFLAG-Halo-WT MDC1 in U2OS ΔMDC1 cells in interphase. Images acquired at 198 frames per second using a Hamamatsu ORCA BT Fusion camera.

**Movie S8.** Live-cell single-molecule imaging movie of transiently expressed 3xFLAG-Halo-WT MDC1 ΔPST1-2 in U2OS ΔMDC1 cells in interphase. Images acquired at 198 frames per second using a Hamamatsu ORCA BT Fusion camera.

**Movie S9.** Live-cell single-molecule imaging movie of transiently expressed 3xFLAG-Halo-WT MDC1 ΔPST1-4 in U2OS ΔMDC1 cells in interphase. Images acquired at 198 frames per second using a Hamamatsu ORCA BT Fusion camera.

**Movie S10.** Live-cell single-molecule imaging movie of transiently expressed 3xFLAG-Halo-WT MDC1 ΔPST1-6 in U2OS ΔMDC1 cells in interphase. Images acquired at 198 frames per second using a Hamamatsu ORCA BT Fusion camera.

**Movie S11.** Live-cell single-molecule imaging movie of transiently expressed 3xFLAG-Halo-WT MDC1 ΔPST1-8 in U2OS ΔMDC1 cells in interphase. Images acquired at 198 frames per second using a Hamamatsu ORCA BT Fusion camera.

**Movie S12.** Live-cell single-molecule imaging movie of transiently expressed 3xFLAG-Halo-WT MDC1 ΔPST1-10 in U2OS ΔMDC1 cells in interphase. Images acquired at 198 frames per second using a Hamamatsu ORCA BT Fusion camera.

**Movie S13.** Live-cell single-molecule imaging movie of transiently expressed 3xFLAG-Halo-WT MDC1 ΔPST1-12 in U2OS ΔMDC1 cells in interphase. Images acquired at 198 frames per second using a Hamamatsu ORCA BT Fusion camera.

**Movie S14.** Live-cell single-molecule imaging movie of transiently expressed 3xFLAG-Halo-WT MDC1 ΔPST in U2OS ΔMDC1 cells in interphase. Images acquired at 198 frames per second using a Hamamatsu ORCA BT Fusion camera.

**Movie S15.** Live-cell single-molecule imaging movie of transiently expressed 3xFLAG-Halo-WT MDC1 in U2OS ΔMDC1 cells in interphase. Images acquired at 147 frames per second.

**Movie S16.** Live-cell single-molecule imaging movie of transiently expressed 3xFLAG-Halo-ΔPST MDC1 in U2OS ΔMDC1 cells in interphase. Images acquired at 147 frames per second.

**Movie S17.** Live-cell single-molecule imaging movie of transiently expressed 3xFLAG-Halo-T>A MDC1 in U2OS ΔMDC1 cells in interphase. Images acquired at 147 frames per second.

**Movie S18.** Live-cell single-molecule imaging movie of transiently expressed 3xFLAG-Halo-T>D MDC1 in U2OS ΔMDC1 cells in interphase. Images acquired at 147 frames per second.

**Movie S19-21.** Laser micro-irradiation of mitotic cells expressing (Movie S19) wildtype Halo-MDC1 or (Movie S20-21) Halo-MDC1 ΔPST (JFX650) from its endogenous locus. Cells were pre-sensitized with Hoechst and images were acquired every second.

**Movie S22.** Live cell timelapse imaging of U2OS ΔMDC1 cells stably expressing HaloTagged wildtype MDC1 (JFX650). Images were acquired every 5 minutes.

**Movie S23.** Live cell timelapse imaging of U2OS ΔMDC1 cells stably expressing HaloTagged MDC1 ΔPST (JFX650). Images were acquired every 5 minutes.

**Movie S24-25.** Live cell timelapse imaging of U2OS ΔMDC1 cells stably expressing HaloTagged MDC1 T>D (JFX650). Images were acquired every 5 minutes.

**Movie S26-28.** Live cell timelapse imaging of U2OS ΔMDC1 cells stably expressing HaloTagged MDC1 T>A (JFX650). Images were acquired every 5 minutes.

**Movie S29.** Live cell imaging of U2OS cells containing a single LacO array expressing wildtype LacI-PST repeat-GFP (left). Cell morphology was detected using transmitted light (middle) and merged with the fluorescence signal (right). Images were acquired every 3 minutes.

**Movie S30-33.** Live cell imaging of U2OS cells containing a single LacO array expressing T>A LacI-PST repeat-GFP (left). Cell morphology was detected using transmitted light (middle) and merged with the fluorescence signal (right). Images were acquired every 3 minutes.

**Movie S34.** Laser micro-irradiation of interphase U2OS ΔMDC1 cells stably expressing wildtype Halo-MDC1 (JFX650, left) and ubiquitin reader 1.0-eGFP (right). Images were acquired every 5 seconds.

**Movie S35.** Laser micro-irradiation of interphase U2OS ΔMDC1 cells stably expressing Halo-MDC1 ΔPST (JFX650, left) and ubiquitin reader 1.0-eGFP (right). Images were acquired every 5 seconds.

**Movie S36.** Laser micro-irradiation of interphase U2OS ΔMDC1 cells stably expressing Halo-MDC1 ΔBRCT (JFX650, left) and ubiquitin reader 1.0-eGFP (right). Images were acquired every 5 seconds.

**Movie S37.** Laser micro-irradiation of interphase U2OS ΔMDC1 cells stably expressing ubiquitin reader 1.0-eGFP. Images were acquired every 5 seconds.

